# Neuropathologically validated MRI to tau PET synthesis via Covariate-modulated attention networks

**DOI:** 10.64898/2025.12.27.696668

**Authors:** Marcell Borhi, Rita Qiuran Lyu, William J Jagust, Jingshen Wang, Joseph Giorgio, Alzheimer’s Disease Neuroimaging Initiative

**Affiliations:** Division of Biostatistics, School of Public Health, University of California, Berkeley, California, USA; Department of Neuroscience, University of California, Berkeley, California, USA, 94720; School of Psychological Sciences, College of Engineering, Science and the Environment, University of Newcastle, Newcastle, New South Wales, Australia, 2308

## Abstract

Tau PET is a powerful tool to assess tau pathology in vivo; however, in comparison to MRI, its development is more recent, is rarely available at scale, and substantially more difficult to acquire. Here, we present Covariate-Modulated Attention UNet (CoMA-UNet) to synthesize subject-specific 3D tau PET from T1 MRI while incorporating in the synthesis procedure readily available covariates. Across six external validation datasets, CoMA-UNet reproduced regional patterns of tau PET uptake showing strong agreement with true PET that was generalizable across tracers. Next, we submitted the synthetic tau PET to a series of downstream clinically relevant tasks. First, MMSE associations between the synthetic tau PET were statistically indistinguishable from true PET. Second, the synthetic tau PET achieved out-of-sample diagnostic classification of dementia with an AUROC=0.99. Third, in two independent autopsy cohorts, voxel wise synthetic tau PET images closely followed neuropathologically defined Braak-stages. Fourth, when fine-tuned on a small subset of atypical presentations of AD, CoMA-UNet preserved phenotype-relevant tau topographies and recovered clinically meaningful structure across atypical clinical phenotypes. These findings demonstrate that the novel CoMA-UNet MRI-based synthesis augmented with covariate information can approximate tau PET with sufficient accuracy for downstream scientific and clinical applications.

## Introduction

Alzheimer’s disease (AD) is a progressive neurodegenerative disorder and the leading cause of dementia worldwide^1,2^ characterized by progressive loss of cognitive and functional abilities^3^. The neuropathological hallmarks of AD are extracellular β-amyloid (Aβ) plaques and intracellular neurofibrillary tangles composed of hyperphosphorylated tau, with the latter showing the tightest coupling to symptom severity, atrophy, and clinical progression^4–6^. To capture these neuropathological hallmarks PET imaging agents have been developed to measure both Aβ and tau in-vivo^7^. Despite the uptake of these imaging technologies in recent years they are relatively new in the research field, with tau-PET imaging only available for about a decade. Accordingly, to leverage the decades of AD biobanking and clinical trials pre-dating tau-PET, novel methods are required to infer the presence and severity of tau pathology in legacy data.

In typical sporadic late onset AD, tau aggregates emerge and spread in a stereotyped sequence captured by the Braak staging framework, beginning in transentorhinal/entorhinal cortex (I/II), advancing to limbic structures (III/IV), and culminating in neocortical association areas (V/VI)^8^. PET-based adaptations of Braak staging recapitulate this topography in vivo, enabling quantitative modeling of disease severity and progression from living patients^9^. Modern biological definitions of AD are established by amyloid (A), tau (T) and neurodegeneration (N) in an ATN framework^3^. Among tau biomarkers, PET imaging with radiotracers such as [^18^F]-flortaucipir (AV-1451), [^18^F]-MK-6240, and [^18^F]-PI-2620 provide unique whole-brain maps of regional tau burden, resolving voxel-level spatial patterns aligned with Braak stages and with strong links to cognition^10–12^. While CSF and plasma phosphorylated tau (e.g., p-tau217) are increasingly accurate for detecting abnormal Aβ and tau and for forecasting decline^13,14^, fluid biomarkers lack spatial specificity; by contrast, tau PET yields topographic information critical for image-based analyses, staging, and anatomically informed treatment monitoring^9^. Despite these strengths, tau PET remains costly and logistically complex due to challenges related to radiochemistry and limited scanner access. As such, tau-PET is sparsely available compared to MRI, thus limiting routine clinical use and leaving tau unmeasured for many individuals and many historical datasets.

One potential way to infer underlying tau-PET levels when it is not available is through the use of tau-PET surrogates. Existing surrogate methodologies typically predict regional summary values rather than full voxel-wise volumes^15–17^; limiting voxel-wise analyses, aggregation with existing volumetric tau-PET data, and downstream applications that depend on spatial fidelity. Recently, volumetric synthesis of tau PET using MRI has been attempted^18^ however these previous approaches have disregarded critical disease related covariate information in the synthesis problem. To address these previous limitations, novel approaches are required to seamlessly integrate predictive covariate information in the synthesis of tau-PET volumetric data from MRI. Furthermore, given the variable access to covariate information (e.g. nascent plasma markers for tau pathology or genetics) this novel formulation must flexibly handle covariate sets with varied missingness.

Here, we present CoMA-UNet, a covariate-conditioned deep learning model for MRI-to-tau synthesis that yields anatomically consistent 3D tau PET volumes in native space. In contrast to regional or tabular surrogates, this voxel-wise formulation aims to preserve disease-relevant topology while remaining compatible with real-world clinical and legacy data constraints, thus maintaining flexibility in downstream tasks. We evaluate the fidelity of the synthetic tau-PET across diverse datasets and tracers. In particular, we highlight the fidelity of the image based on ground truth neuropathological staging and examine image utility in a range of downstream tasks including re-stratification of clinical trials based on anticipated levels of baseline tau pathology. Together, we present CoMA-UNet as an effective method to assess in-vivo tau pathology when tau PET is missing.

## Results

### Participants and Datasets

Analyses were performed across seven discrete datasets derived from three parent cohorts (ADNI, A4, and NACC), including cross-validation, independent held-out cross-sectional, cross-tracer validation, neuropathology-linked evaluations, and atypical presentation performance. Training and internal cross-validation cohorts were drawn from the Alzheimer’s Disease Neuroimaging Initiative (ADNI) and the Longitudinal Evaluation of Amyloid Risk and Neurodegeneration (LEARN) Study subset of the Anti-Amyloid Treatment in Asymptomatic Alzheimer’s (A4) Study (ADNI-A4 training; n=1250 MRI-tau PET image pairs; 684 clinically unimpaired, 228 clinically impaired). These participants had paired T1-weighted MRI and [^18^F]flortaucipir (FTP) tau PET imaging as well as thirteen covariates grouped into demographic (age, sex, education, marital status, race, height, weight), vital sign (systolic and diastolic blood pressure, pulse, respiratory rate, temperature), genetic (APOE4 allele count) features, and where available, plasma biomarkers (p-tau217, NfL, GFAP, Aβ40, and Aβ42). Validation within the ADNI-A4 training set was performed using five-fold cross-validation, with all presented results from the held-out test folds. Among external validation datasets, varying degrees of overlapping MRI and co-variate or ground truth tau PET data was available (Figure 1). The external validation datasets included (a) 100 ADNI and 100 A4 held out participants (cross-sectional validation; n=200; 81.5% with plasma ptau-217 available), (b) The National Alzheimer’s Coordinating Center (NACC) Standardized Centralized Alzheimer’s & Related Dementias Neuroimaging (SCAN) dataset (NACC-SCAN; n=1291 subjects, 1409 observations, with 0% plasma ptau-217 available), (c) Treatment and placebo arms from the A4 clinical trial excluding the participants from the A4-LEARN present in the ADNI-A4 training data (A4 validation; n = 795, with 91.2% plasma ptau-217 available), and two datasets with paired neuropathology assessment, (d) ADNI with neuropathology (ADNI-NP validation; n=99, with 99% available ptau-217), and (e) NACC participants with non-SCAN-compliant MR imaging (NACC non-SCAN; n=475, with 0% plasma ptau-217) (**Error! Reference source not found.**). Results reported in the external validation datasets (b-e) are generated from the fixed model trained on the complete ADNI-A4 training dataset.

**Figure 1.**
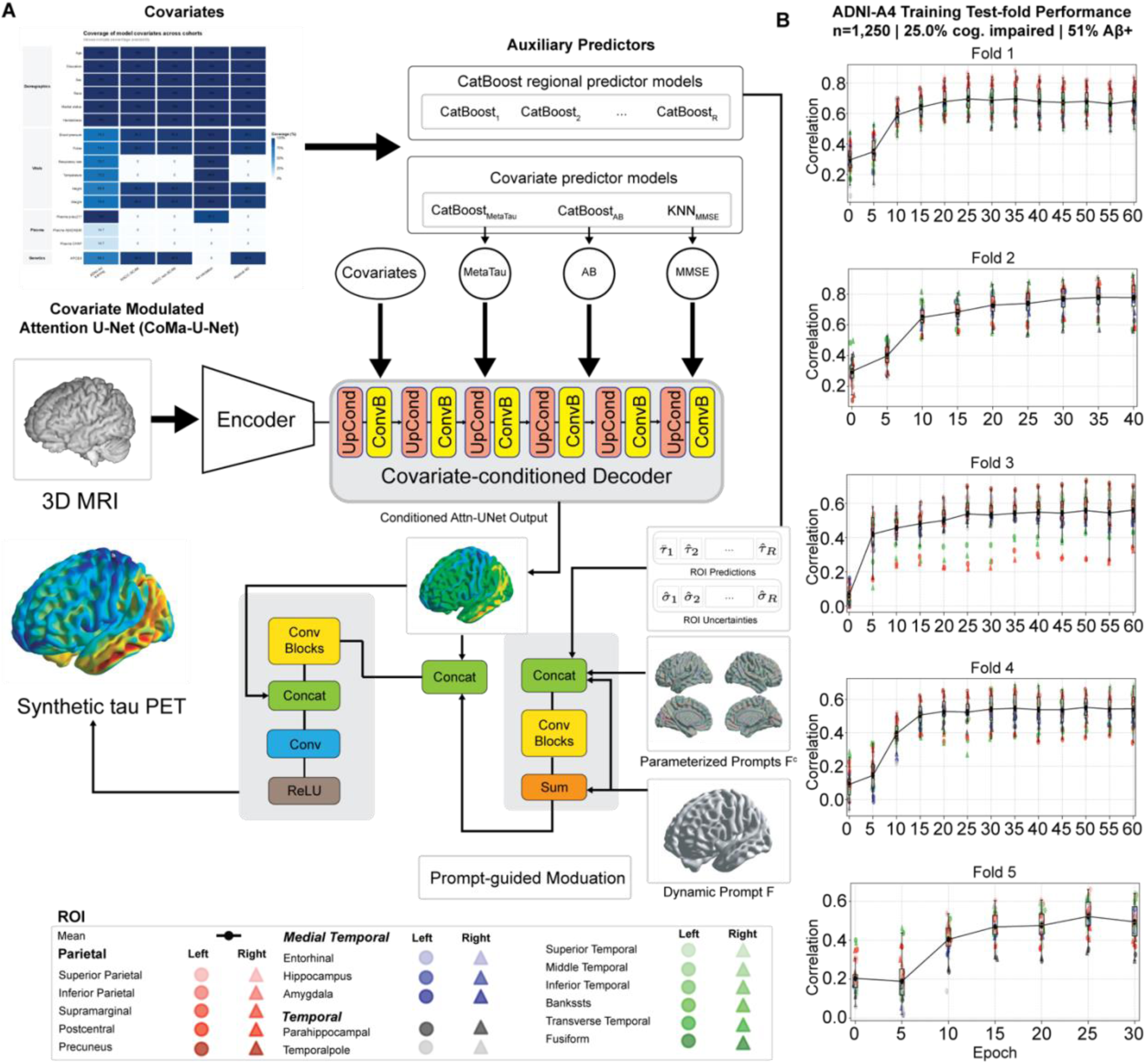
(A) Overview of the Covariate-Modulated Attention U-Net (CoMA-UNet) architecture. A 3D MRI is encoded through a convolutional encoder, while subject-level covariates (vital, demographic, MRI-derived features, genetic variables, and plasma biomarkers) are transformed into dynamic, parameterized prompts. These prompts condition the decoder via concatenation and covariate-modulated attention blocks, enabling voxel-wise tau prediction that adapts to individual covariate profiles. The model outputs a full synthetic FTP tau PET volume. (B) Training curves for all five folds of cross-validation within the ADNI-A4 training dataset, showing correlation between predicted and ground-truth regional FTP SUVR values in the held-out test partition at each test epoch. Colored points correspond to individual regions of interest (ROIs), while the black line indicates the average correlation across ROIs (CorrAVG). Across all folds, model performance steadily improves with training, with both CorrAVG and ROI-specific correlations showing consistent convergence.

Finally, to incorporate atypical clinical presentations of AD in model development, we included a separate subset of NACC participants with atypical clinical phenotypes and available SCAN-compliant imaging (Atypical SCAN; n = 253; Table 2). As atypical presentations were not represented in the ADNI-A4 training cohort, these data were used to fine tune the ADNI-A4-trained model. Accordingly, performance was evaluated using a separate five-fold cross-validation strategy within the atypical SCAN sample. To test model generalizability, we included as a held-out-sample NACC participants with atypical presentations and non-SCAN-compliant MR imaging (Atypical non-SCAN n = 199; Table 2). To ensure the fine-trained model did not forget mapping for typical AD participants we tested the fine-tuned model on a held-out-sample of participants with typical presentations of AD and corresponding ground truth tau PET (Typical SCAN n = 99; Table 2).

**Table 1.**
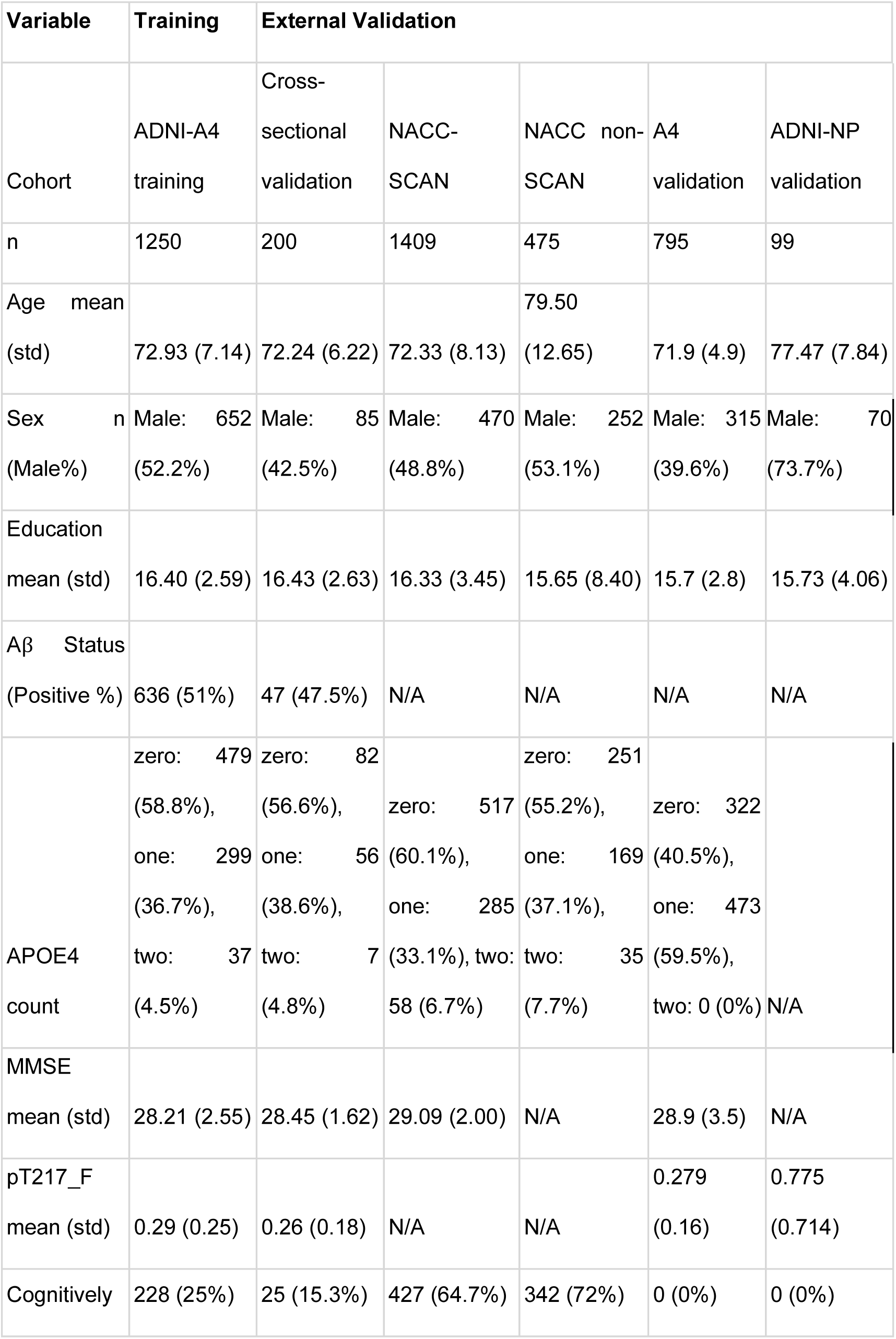
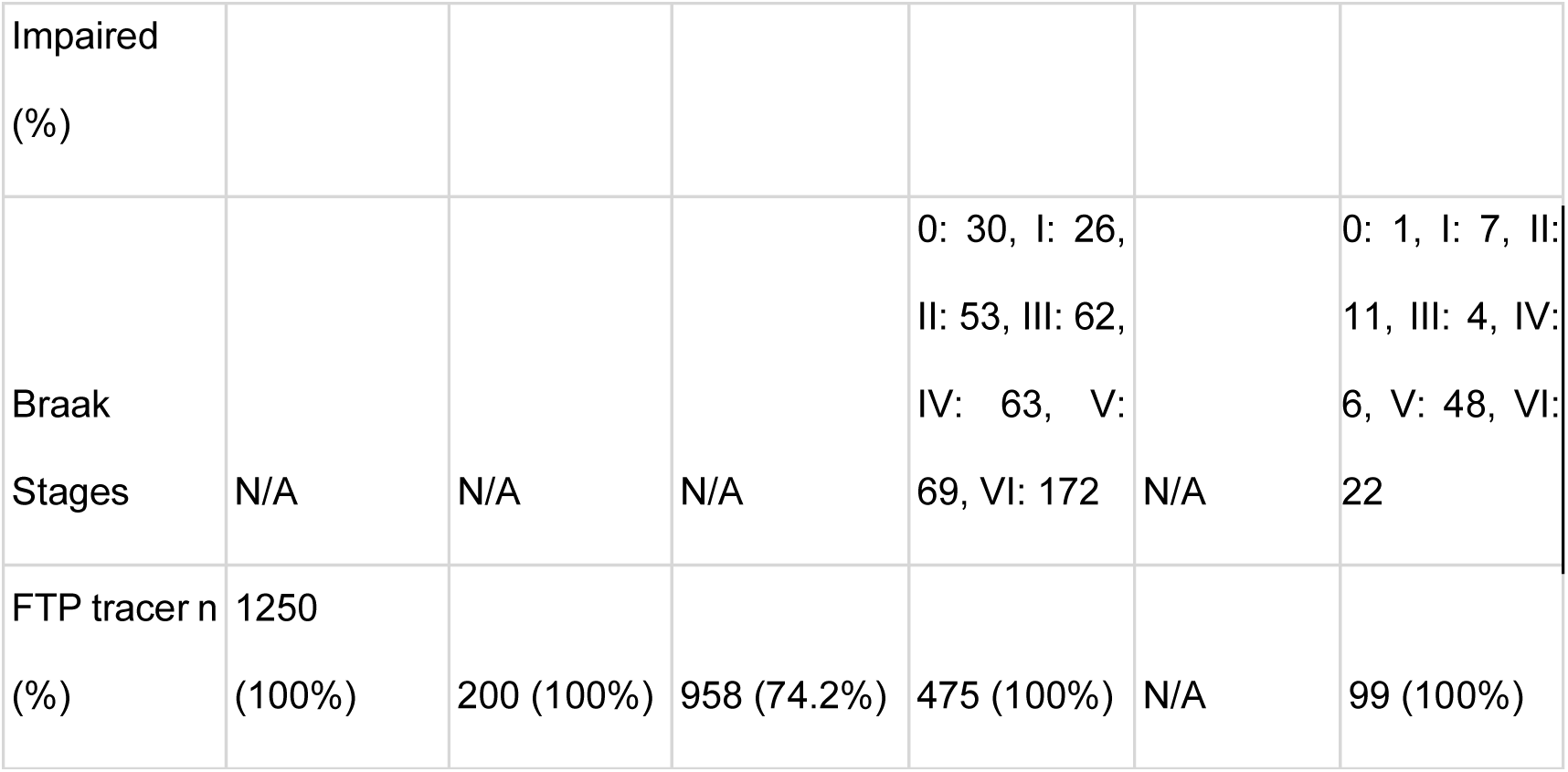
Cohort-level demographics, clinical variables, and imaging availability for all datasets included in this study. Values represent means (±SD) for continuous variables and counts (percent) for categorical variables.

**Table 2.** Summary characteristics are shown for atypical SCAN, atypical non-SCAN, and typical SCAN participants, including age, sex, education, APOE4 category counts, and dementia prevalence. Continuous variables are reported as mean (SD), and categorical variables are reported as n (%).

| Dataset | n | Age mean (std) | Sex n (Male%) | Education mean (std) | APOE4 count | Dementia (%) |
| --- | --- | --- | --- | --- | --- | --- |
| Atypical SCAN | 253 | 71.1 (8.5) | 124 (47.5%) | 16.2 (2.6) | zero: 117, one: 51, two: 60 | 170 (65.1%) |
| Atypical non-SCAN | 199 | 76.3 (10.6) | 106 (53.3%) | 16.5 (8.8) | zero: 107, one: 66, two: 24 | 130 (65.3%) |
| Typical SCAN | 99 | 72.5 (9.5) | 41 (41.4%) | 16.6 (2.7) | zero: 20,<br>one: 24,<br>two: 15 | 98 (99%) |

### Cross-validation and Cross-sectional Performance of Covariate-modulated Attention UNet

CoMA-UNet was built to incorporate AD related covariates into the voxel-to-voxel Attention-UNet in three stages. First, covariates are used in a series of CatBoost models estimating regional tau burden, temporal meta-ROI tau (MetaTempTau), Aβ positivity, and MMSE. In the second stage, the deep learning model integrates predicted values of MetaTempTau, Aβ PET status, and MMSE estimates through a conditional decoder. In the final stage, regional predicted tau estimates are used to modulate the decoded image from the conditioned attention-UNet output. In doing so, disease related covariate information is used to conditionally up sample based on predicted AD severity (i.e. MetaTempTau, Aβ, and cognitive impairment), while also embedding regionally specific covariate related signal thorough the spatially constrained modulation of the up sampled image (Figure 1A). No PET or cognitive variables are included in the image synthesis at inference, with images generated using the T1-weighted MRI and any available covariates from the selected categories (demographic, vital sign, genetic, and plasma biomarkers). Critically, prior to running the CatBoost prediction model an initial imputation model is run to fill in missing covariate data. In this way, CoMA-UNet seamlessly handles variable missingness in the covariate feature set allowing the model to synthesise tau PET images with any, all, or no covariate data available. We present findings in discrete samples with varied degrees of covariates available testing the utility and reliability of synthetic tau PET generated with or without plasma markers (Figure 1A).

Synthetic tau PET images were generated using the CoMA-UNet framework trained and internally validated across five-folds of the ADNI-A4 dataset (n=1250). Performance metrics include the average correlation (Pearson’s) coefficient (Corr_AVG_) of predicted and ground truth tau in selected regions of interest (ROIs) (Supplementary Table 1), average voxel-wise mean absolute percentage error (MAPE) and mean absolute error (MAE), and the structural similarity index (SSIM). Across the five folds, the converged models achieved an average held out prediction performance of Corr_AVG_=0.5180 (± 0.034), MAPE =10.88% (± 1.23), MAE =0.143 (± 0.021), and a SSIM=0.823 (± 0.017) (Figure 1B). Fidelity was assessed across all held-out cross-validation folds in the ADNI-A4 training dataset by correlating synthetic and actual tau PET SUVRs in key ROIs (Supplementary Figure 1). In key AD related regions, a high cross-validated correlation was achieved in inferior temporal (r = 0.63), middle temporal (r = 0.63), as well as key Parietal lobe regions, including the inferior parietal (r = 0.64) and precuneus (r = 0.63). The trained model was then applied to the fully held out cross-sectional validation dataset (n = 200). CoMA-UNet attained consistent performance, with Corr_AVG_ = 0.462, MAPE = 10.87%, MAE = 0.131, and SSIM=0.79 (Supplementary Figure 2).

We next contrasted the goodness-of-fit results of CoMA-UNet to alternate model architectures, including Attention UNet, UNet, UNETR, and UNETR variant with attention gates in the Decoder (Supplementary Table 2). The CoMA-UNet produced substantial gains over all alternative architectures, which typically achieved Corr_AVG_ < 0.15 across the five folds. The Attention UNETR, and Attention UNET models had the next lowest average regional MAPEs with 12.96% (±2.21), and 13.05% (±1.43) in the cross-validation, respectively. The SSIM of all architectures were relatively high, with the highest average SSIM being 0.7506 achieved by the Attention UNet.

### Performance on SCAN-compliant MRIs from the NACC-SCAN dataset & generalizability to different radiotracers

We next used the NACC-SCAN dataset to assess the model’s ability to synthesize tau PET without the inclusion of plasma ptau-217 as a covariate. This dataset consisted of entirely held out subjects from the NACC cohort with SCAN compliant tau PET (n=1409), and included a mix of tau PET tracers (n=958 FTP tau PET; n=451 MK-6240 tau PET), allowing us to also assess if the model could capture variance in tau PET images acquired with an unseen radiotracer. Since CoMA-UNet was trained to synthesize FTP tau PET images that operate on a different scale from MK tau PET, we assessed predictive performance using Pearson correlation, excluding other metrics that are sensitive to scale differences.

We observed strong performance in both Pearson correlation and *R*^2^ with both the FTP and MK-6240 tau PET (Supplementary Figure 3). In particular, the CoMA-UNet model achieved correlations of r=0.471 in the entorhinal, and an overall average of r=0.486 in the temporal regions. Across both tracers, the model attained a Corr_AVG_ of r=0.498. Despite not being trained on images with the MK-6240 radiotracer CoMA-UNet maintained similar performance, indicating that it can reproduce the underlying association between MRI and tau PET signal. Notably, the model achieves these results in the absence of plasma biomarkers, underpinning the predictive capabilities of the model under different patient populations, even with few available covariates.

### Classification of CN, MCI and Dementia using Synthetic tau PET

We next evaluated whether the average synthetic tau PET signal in the meta temporal ROI (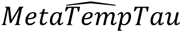) generated by CoMA-UNet showed diagnostic utility. As all samples drawn from the A4 datasets had clinical diagnoses of CN, we limited this analysis to ADNI samples (n = 602; 7% with plasma ptau-217), avoiding any biases from population differences.

Within the five-folds of the ADNI-A4 training dataset, we evaluated diagnostic classification using only completely held-out (ADNI) subjects from each fold. For each fold, a logistic-regression classifier was trained on synthetic 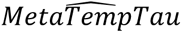 values from the remaining four folds and evaluated on that fold’s held-out participants. Across the five folds, CN vs Dem classification achieved ROC-AUC values exceeding 0.80 in four of the five held-out subsets (Figure 2B). We further evaluated the discriminative utility of the synthetic tau PETs in the completely held-out cross-sectional validation samples. To protect against bias resulting from population differences, we again only considered classification in the ADNI subjects (n= 63; 100% with plasma ptau-217). Similar to the cross-validation findings, the logistic regression classifier using 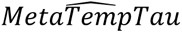 in the cross-sectional validation data set showed strong discriminative power (AUROC=0.99) in the CN vs Dem classification (Figure 2D).

**Figure 2.**
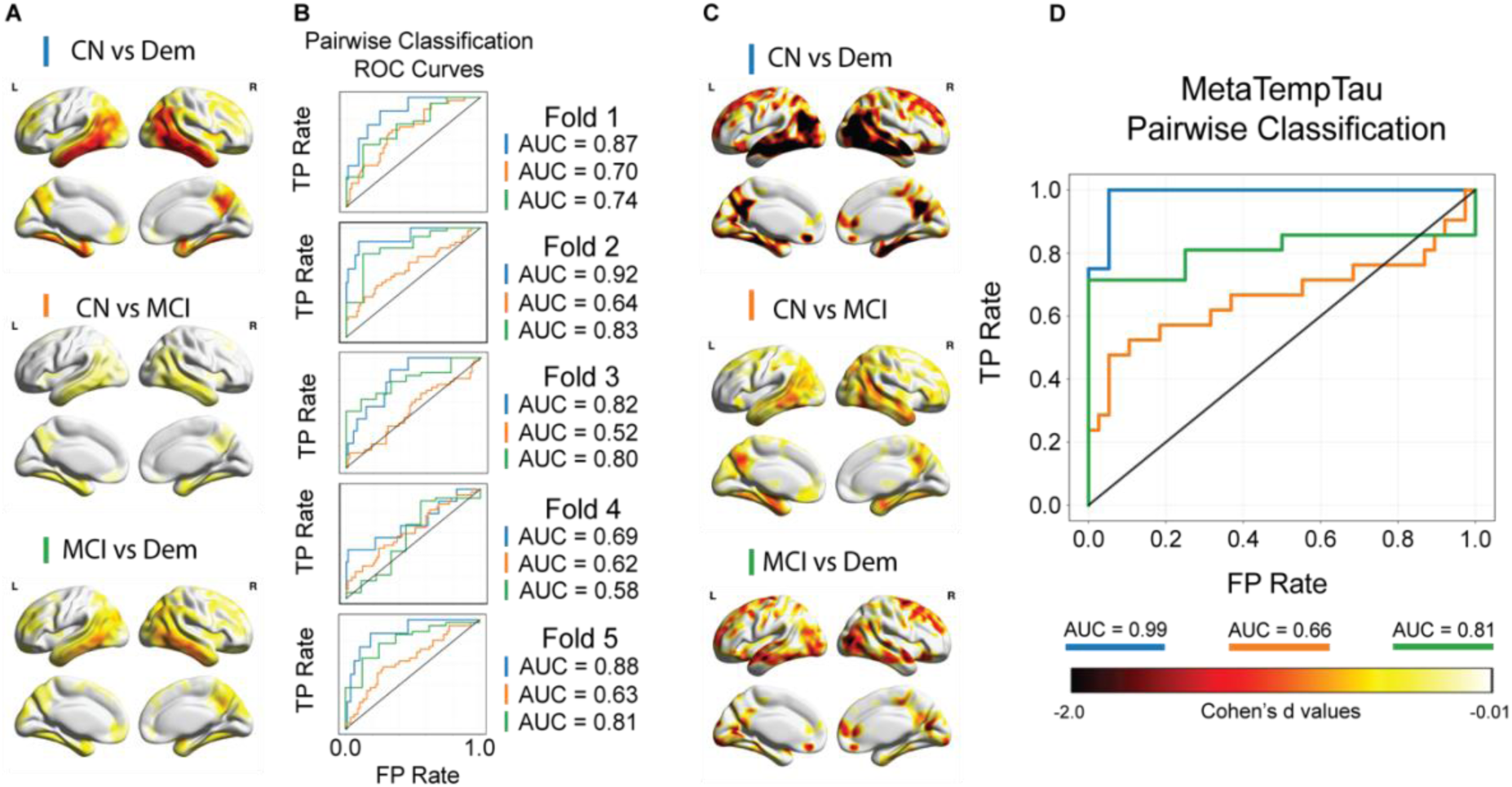
(A) Voxel-wise Cohen’s *d* values within all held-out cross-validation samples in the ADNI-A4 training dataset show large effect sizes (*d* < −0.8) in the temporal regions for all comparisons. (B) ROC Curves for logistic regression classifiers using 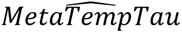 across each of the held-out samples across the five folds of the ADNI-A4 training dataset, showing consistent discriminative performance. (C) Voxel-wise Cohen’s *d* values for each pairwise classification within all held-out cross-sectional samples show very large effect sizes (*d* < −2) in the Temporal and Parietal regions for all comparisons. (D) Receiver operator curve for a logistic regression classifier using synthetic 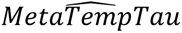 in the CN vs Dem, and MCI vs Dem pairwise classification tasks.

Finally, we evaluated the discriminative utility of the synthetic tau PET on the voxel level running voxel-wise two sample t-tests to assess mean difference between levels of clinical impairment. As before we excluded A4-LEARN subjects using the held-out ADNI synthetic tau PET volumes generated from the five-fold cross-validation procedure. We computed unthresholded voxel-wise Cohen’s *d* in the MNI-space normalized synthetic tau PETs for both the held out ADNI training dataset samples (Figure 2A) and cross-sectional validation samples (Figure 2C). Across both cohorts, we observed particularly large effect sizes (*d* < −2) in the Temporal and Parietal lobes for the CN vs Dem comparison, with more modest effect sizes for CN vs MCI and MCI vs DEM comparisons.

### Cross sectional MMSE prediction

We next examined the regional association between synthetic tau PET and MMSE, and MRI thickness and MMSE in the cross-sectional validation dataset (n = 198 with available MMSE, 63.1% with plasma ptau-217) to assess how well synthetic tau PETs capture subject-wise levels of cognition. We then contrasted the differences in these associations between imaging measures and MMSE using Steiger’s Z tests. This revealed that synthetic tau PET values had a stronger association with cognition in the temporal and parietal lobe regions than the MRI used to generate the synthetic tau PET images. When analyses were stratified by Aβ PET positivity status, the synthetic tau PETs within the Aβ positive subgroup showed significantly stronger association with subject baseline MMSE scores than MRI volume (Benjamini–Hochberg correction, *q* < 0.05, *p*_FDR_ = 0.0239). This was observed in tau-PET AD related regions such as the banks of the superior temporal sulcus (*Z* = -2.581, *p*_uncorrected_ = 0.0049 < 0.0239), inferior parietal (*Z* = -2.730, *p*_uncorrected_ = 0.0031 < 0.0239), middle temporal (*Z* = -2.823, p_uncorrected_ = 0.0023 < 0.0239), and precuneus (-2.443, *p*_uncorrected_ = 0.0072 < 0.0239) (Figure 3A). At the voxel-level in spatially normalized MNI space, we observed strong correlations in the temporoparietal regions (*r* < -0.5), with weaker correlation in the frontal regions (Figure 3B) in Aβ positive individuals. Within the Aβ negative subgroup however, there were no significant differences in the association between MRI and MMSE and synthetic tau PET and MMSE, suggesting that the synthetic tau PET captured AD related variance in cognition (i.e. dependent on Aβ status). Finally, we found that within the cross-sectional validation dataset, the association between the 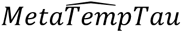 and MMSE (*r* = -0.442, df = 196, 95% CI [-0.55, -0.32], *p* < 0.001; Figure 3D) did not differ significantly (*Z* = 0.974, df = 195, *p* = 0.1650) from the ground truth association between *MetaTempTau* and MMSE (*r* = - 0.491, df = 196, 95% CI [-0.59 , -0.38], *p* < 0.001; Figure 3C). Indicating that synthetic tau PET captures individual variability in MMSE comparable to ground truth tau PET.

**Figure 3.**
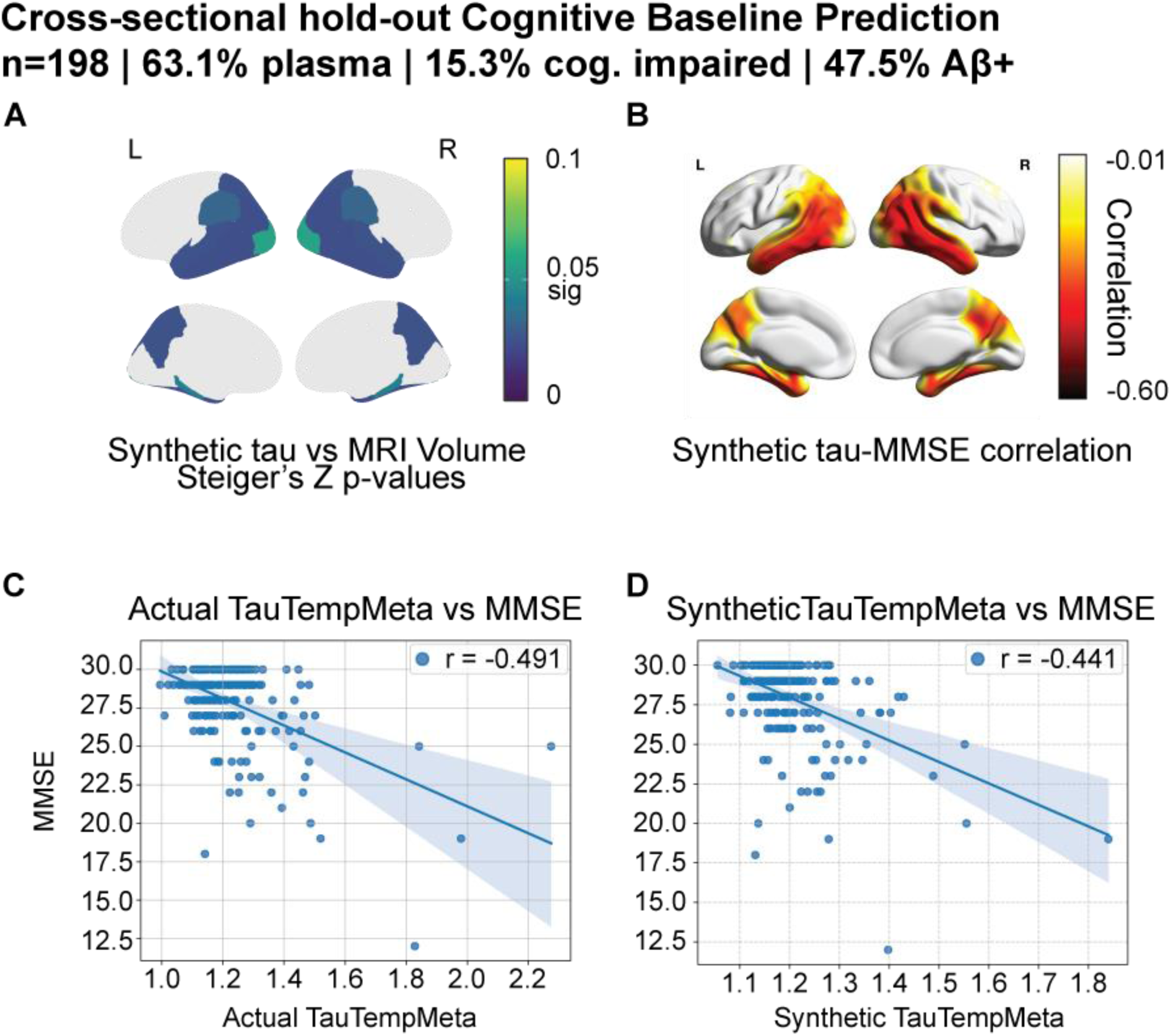
(A) Benjamini & Hochberg Family-wise Error corrected p-values for Steiger’s Z values between synthetic tau PET values and MRI volumetric measures within amyloid positive subpopulation of the cross-sectional validation dataset subjects. (B) Voxel-wise correlation between synthetic tau PET and MMSE among amyloid positive subjects. (C, D) Correlation between the ground-truth *MetaTempTau* and MMSE vs 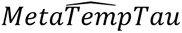 and MMSE.

### Performance on A4 samples with only MRI observed

To evaluate the utility of the model to predict prospective cognitive decline in asymptomatic individuals, we re-stratified participants in the A4 validation dataset (n = 795) into three groups according to tertile thresholds of 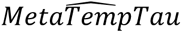 values derived from synthetic tau PET scans. Using these tertile assignments, we computed longitudinal PACC trajectories within the treatment and placebo arm using natural cubic splines consistent with the modeling approach in the A4 study^19^. We repeated this same procedure using synthetic tau PETs generated without plasma ptau-217 as a covariate input to CoMA-UNet, yielding corresponding trajectories for the treatment and placebo groups, thereby probing the robustness of the model to missing covariate information.

Across all conditions, participants in the highest 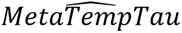 tertile (T3) showed steeper cognitive decline relative to those in the lowest tertile (T1). In the treatment arm, differences in PACC trajectories were significant when using plasma biomarkers in the input (T3–T1 = –3.11 ± 0.64, t = –4.82, p = 1.5 × 10⁻⁶; T3–T2 = –2.42 ± 0.64, t = –3.79, p = 1.5 × 10⁻⁴; Figure 4B). These differences remained significant without plasma biomarker inputs between T3 and T1 ( *t* = -4.82, *p* < 0.0001), as well as between T2 and T3 (*t* = -3.29, p-value = 0.001; Figure 4A). Similarly, in the placebo arm, participants in T3 also exhibited greater cognitive decline both when using plasma inputs (T3–T1 = –2.64 ± 0.47, t = –5.57, p = 2.7 × 10⁻⁸; T3–T2 = –2.37 ± 0.47, t = –5.05, p = 4.6 × 10⁻⁷; Figure 4D), and without plasma (T3–T1 = –1.91 ± 0.47, t = –4.09, p = 4.4 × 10⁻⁵; T3–T2 = –1.74 ± 0.49, t = –3.58, p = 3.4 × 10⁻⁴; Figure 4C).

**Figure 4.**
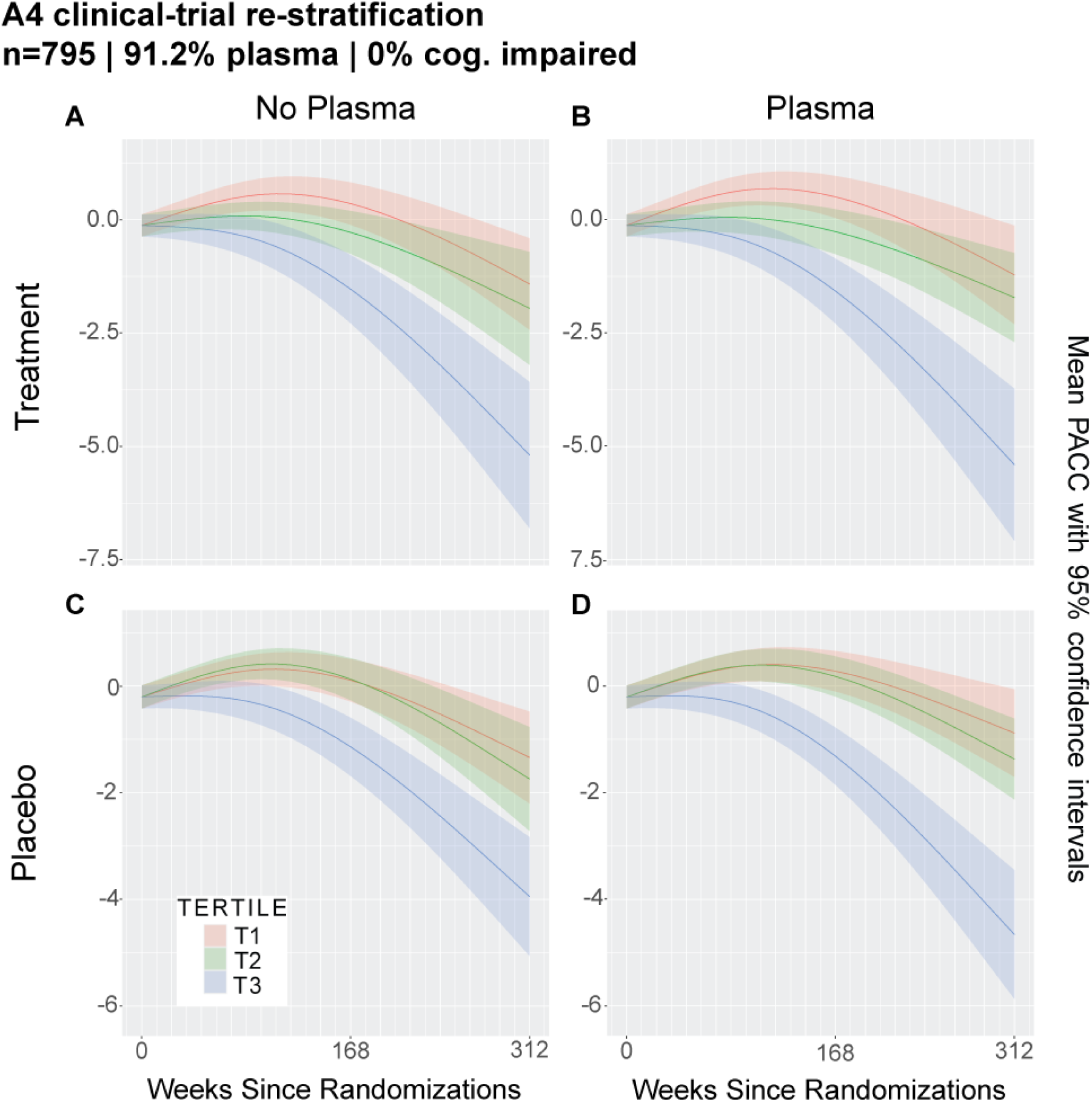
Held-out A4 subjects stratified according to tertiles of average temporal meta ROI SUVR derived from synthetic tau PETs generated by the CoMA-UNet including plasma ptau-217 as a covariate (panels B and D), and in the absence of plasma ptau-217 as a covariate (panels A and C). Panels A and B display PACC score trajectories for each tertile among subjects that received treatment. Panels C and D display PACC score trajectories for each tertile among control (placebo) subjects.

### Synthetic tau PET association with Braak Stage neuropathology

We next assessed how well the synthetic tau PET images derived using the CoMA-UNet recapitulated underlying Braak stage pathology as assessed by post-mortem neuropathology assessment. Using the ADNI-NP validation dataset (n=99; 99% with plasma ptau-217 biomarkers), which consists of subjects with autopsy-confirmed Braak stage data, we evaluated how faithfully the synthetic tau PET derived ante-mortem could track the gold standard neuropathological progression of tau. We grouped subjects by adjacent Braak stages: Braak Stage group I/II, III/IV, and V/VI and contrasted the voxel-wise uptake difference between adjacent tau stages. The analysis revealed a “spread” of tau from medial temporal lobe (stage I/II) to other temporal regions, posterior cingulate/retrosplenial cortex, parietal & frontal regions (stage III/IV), and finally diffuse neocortical regions (stage V/VI) recapitulating the underlying stages of tau neuropathology (Figure 5A, B). Ordinal regression of 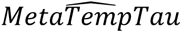 signal against Braak group confirmed these voxel-level findings, showing a monotonic increase in the predicted probability of higher Braak stage groups with increasing temporal tau burden (*β* = 10.69, 95% *CI*[7.25,14.12], *z* = 6.10, *p* < 0.0001) (Figure 5C; Supplementary Figure 4, 5).

**Figure 5.**
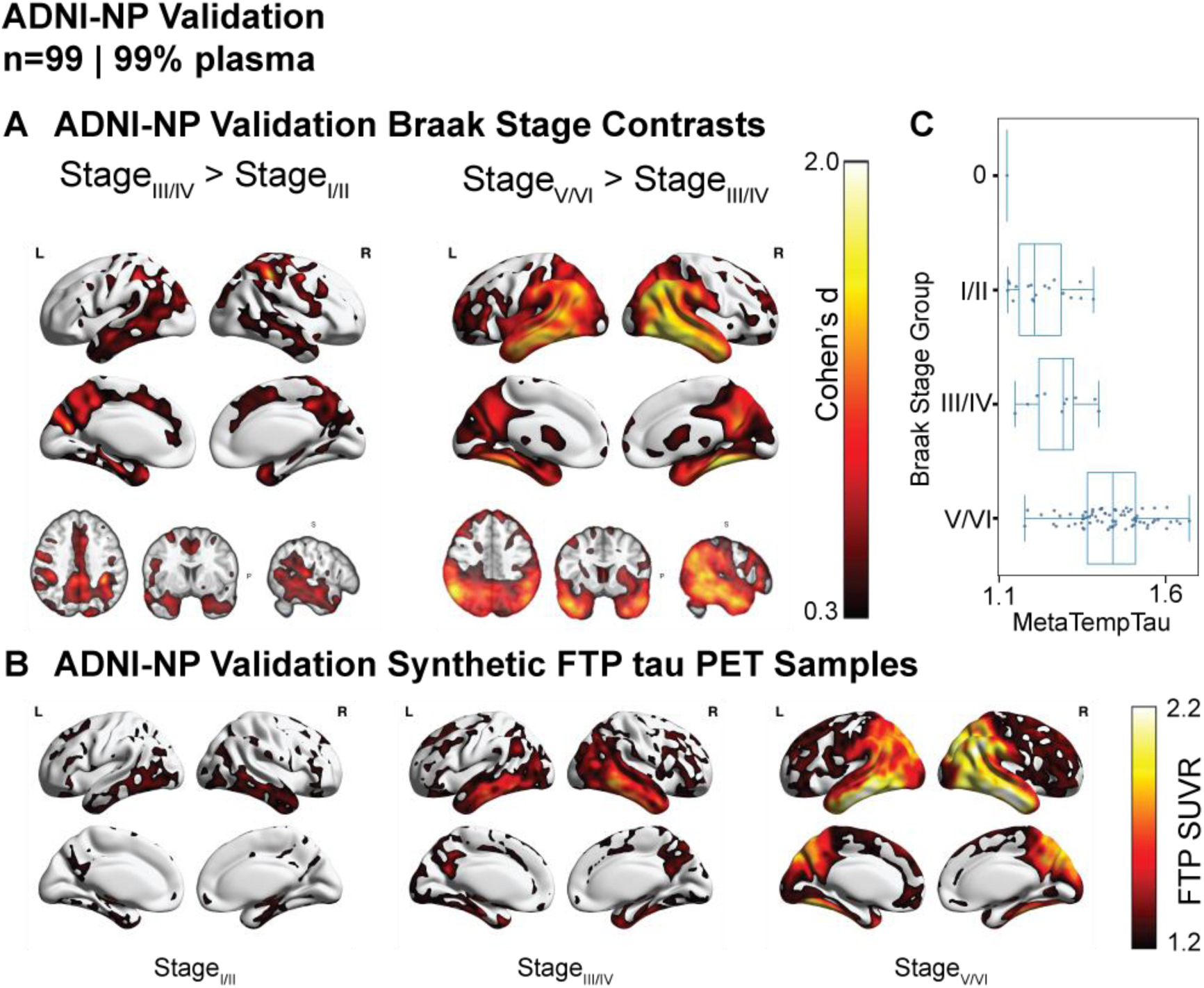
Braak-stage effects in ADNI neuropathology (NP) participants using synthetic tau PET (Braak stage 0/I/II/III/IV/V/VI: n= 1/7/11/4/6/48/22) (A) Voxel-wise Cohen’s *d* maps comparing synthetic tau deposition between Braak stage I/II vs III/IV (left) and III/IV vs V/VI (right). The contrast between I/II and III/IV shows focal effects in medial and inferior temporal regions, whereas the contrast between III/IV and V/VI shows larger clusters across lateral temporal, parietal, and occipital cortex. (B) Representative synthetic tau PET images from Braak Stage I/II, III/IV, and V/VI (left to right), showing low medial-temporal uptake in I/II, higher and more diffuse temporal uptake in III/IV, and the highest SUVR levels with broad cortical involvement in V/VI. (C) Boxplots of average MTL SUVR for each Braak Stage group. Median MTL SUVR increases from low values in I/II, to intermediate levels in III/IV, to highest values in V/VI, matching the uptake differences visible in Panel B.

### Synthetic tau PET generated using non-SCAN compliant MRIs from the NACC dataset

Finally, we assessed if CoMA-UNet could generate high fidelity synthetic tau PET in a heterogeneous sample with non-standardized MRI imaging protocols and no available plasma ptau-217 data. Using the NACC non-SCAN dataset, comprising non-SCAN-compliant MRIs (n = 475) with post-mortem assessment, we generated a synthetic tau PET volume for each subject using the CoMA-UNet. Across adjacent Braak stage contrasts, the synthetic images recapitulated the expected neuropathological progression. Relative to Braak I/II, subjects in Braak III/IV exhibited significantly greater simulated uptake within the medial temporal lobe, within the Braak V/VI group there were further increases extending from the medial temporal lobe into temporal neocortex and association cortices (Figure 6A, B). Effect sizes were largest in temporal regions, with the III/IV versus V/VI comparison revealing a marked neocortical spread consistent with late-stage tau deposition (Figure 6A, B). Complementing the voxel-wise contrasts, an ordinal regression of 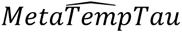 signal against Braak stage demonstrated a monotonic increase in the predicted probability of higher Braak stage groups with increasing temporal tau (*β* = 2.96,95%*CI*[2.10,3.81], *z* = 6.78, *p* < 0.001), indicating that a simple summary of the synthetic images also tracks ordinal pathology (Figure 6C; Supplementary Figure 6, 7).

**Figure 6.**
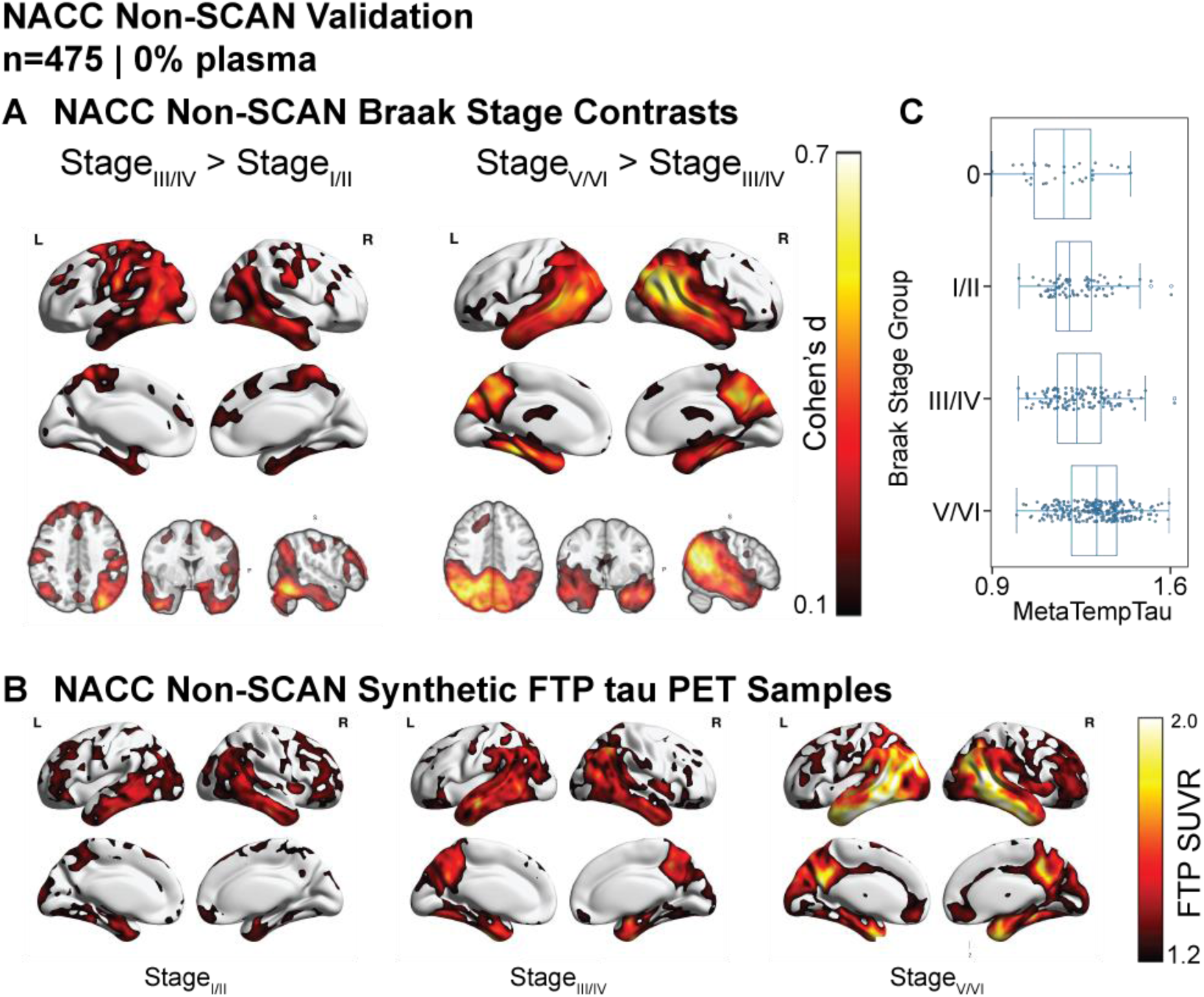
Braak-stage effects in NACC non-SCAN participants using synthetic tau PET (Braak stage 0/I/II/III/IV/V/VI: n = 30/26/53/62/63/69/172). (A) Voxel-wise Cohen’s *d* maps showing regional differences in synthetic tau deposition between Braak Stage I/II vs. III/IV (left) and III/IV vs. V/VI (right). The I/II vs III/IV contrast shows strongest effects in medial temporal regions extending into inferior temporal cortex, whereas the III/IV vs V/VI contrast shows broad neocortical association involvement. (B) Representative synthetic tau PET images from individuals in Braak Stage groups I/II, III/IV, and V/VI (left to right), illustrating the progressive spatial spread of tau. (C) Boxplots of average meta-temporal (MTL) SUVR across Braak Stage groups (I/II, III/IV, V/VI), demonstrating monotonic increases in regional tau signal with advancing Braak stage.

### Synthetic tau PET in atypical presentations of AD

We next evaluated CoMA-UNet in NACC participants with atypical clinical presentations of AD and available SCAN-compliant imaging. For each fold, the model trained on the full ADNI-A4 dataset was fine-tuned on four folds and used to generate synthetic tau PET only for the held-out fold. Held-out predictions were then stacked across folds to obtain out-of-sample synthetic tau PET images for all participants. Downstream analyses in completely held out data from the atypical non-SCAN and typical SCAN cohorts used ensemble-averaged synthetic tau PET volumes, obtained by voxel-wise averaging predictions from the five folds of the model trained on the atypical SCAN dataset.

Across the five folds, the converged models achieved an average held out prediction performance of Corr_AVG_=0.5727 (± 0.0604), MAE = 0.0543 (± 0.003), and a SSIM= 0.819 (± 0.025).

Representative subject-level examples indicate the model was able to capture topographies of tau PET in a phenotypically heterogenous population. This was observed on the single subject level; with synthetic tau PET in a language-predominant atypical AD participant qualitatively reproducing the asymmetric tau distribution observed in ground-truth tau PET with strong asymmetry agreement shown as hemispheric percent-ratio values. In comparison, a typical AD case showed more symmetric uptake and weaker hemispheric-ratio correspondence (Figure *7*). These results indicate that CoMA-UNet can recover regional tau PET signal in an atypical phenotype-enriched sample despite greater clinical heterogeneity than the primary ADNI-A4 training cohort.

**Figure 7.**
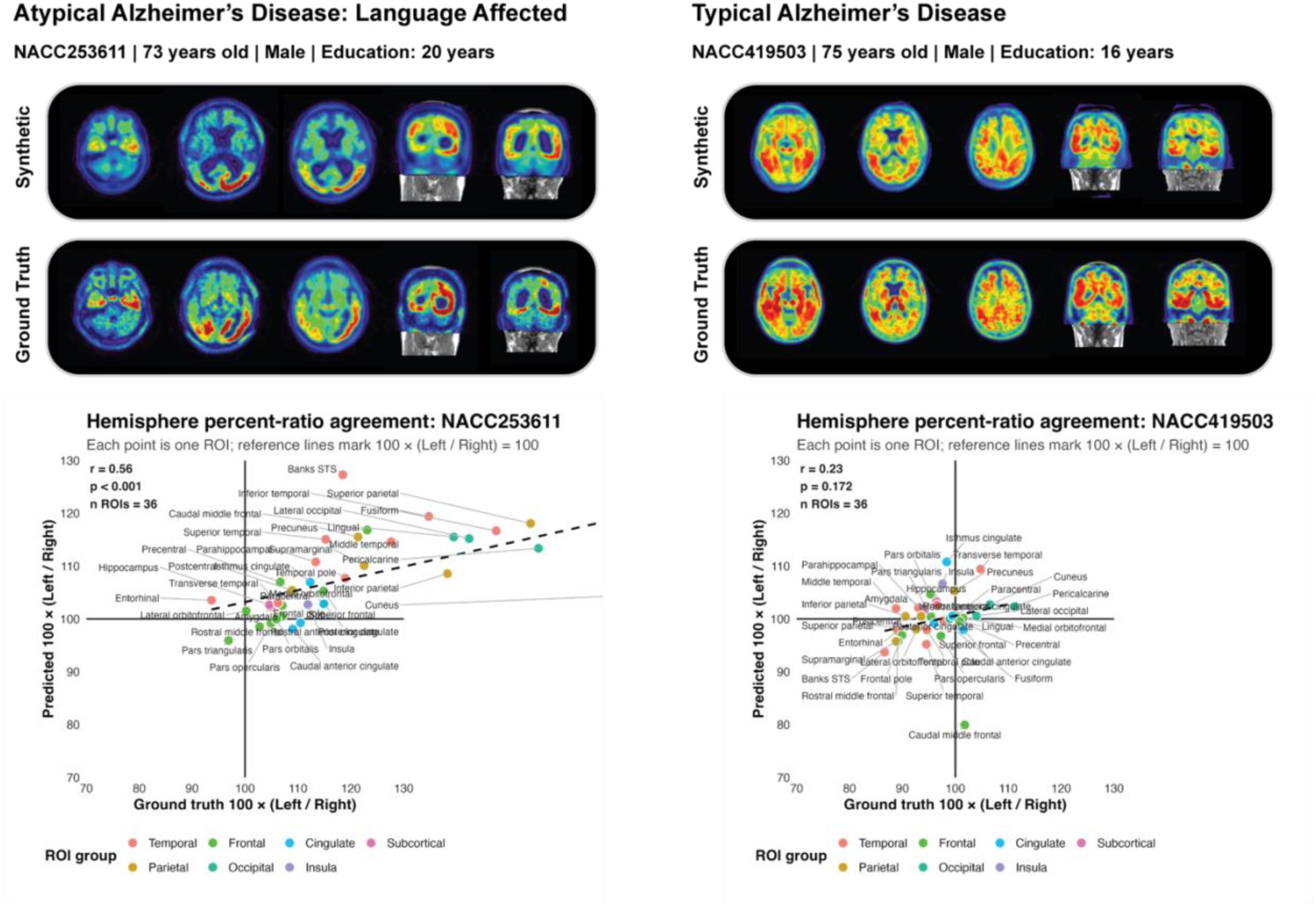
Representative preservation of hemispheric tau PET asymmetry in atypical AD. Representative ground-truth and synthetic tau PET images are shown for a language-predominant atypical AD participant and a typical AD comparison participant from NACC. Scatterplots quantify subject-level agreement between ground-truth and synthetic hemispheric asymmetry across bilateral ROIs. Each point represents one ROI, with hemispheric asymmetry expressed as 100 × (left SUVR / right SUVR); horizontal and vertical reference lines indicate no hemispheric asymmetry (100). In the language-predominant atypical AD case, synthetic tau PET preserved the ground-truth pattern of hemispheric asymmetry across ROIs (r = 0.56, p < 0.001, n = 36 ROIs). In the typical AD comparison case, hemispheric asymmetry was less pronounced and ROI-level percent-ratio agreement was weaker (r = 0.23, p = 0.172, n = 36 ROIs). Points are colored by ROI group.

To test if there was consistent topographical structure between the synthetic and ground truth tau PET in atypical clinical phenotypes we performed k nearest neighbors clustering on the regional values. Running the clustering on the same atypical sample of ground truth tau PET and held-out cross fold synthetic tau PET (i.e. aggregating inference folds not used in the training fold) showed highly similar results. In particular, we observed strong centroid alignment between SCAN ground-truth and SCAN cross-fold synthetic tau PET, with matched correlations of r = 0.810, 0.952, 0.843, and 0.927, corresponding to a mean correlation of r = 0.883 ± 0.065. This correspondence indicates that the dominant topographic structures in ground truth tau PET in atypical phenotypes is highly consistent with the dominant topographic structures in the synthetic tau PET.

We next tested if unsupervised clustering of the synthetic tau PET signal corresponded to phenotypic variation assigned as an atypical AD clinical phenotype in the NACC (Language, Visuospatial, Attention, Executive). Synthetic tau PET was generated on held out atypical non-SCAN participants, using as input heterogenous T1 weighted MRI and no plasma markers. Clustering of non-SCAN predicted tau PET yielded four clusters of sizes 41, 66, 71, and 21, with cluster membership significantly associated with domain label (χ² = 25.24, df = 12, p = 0.014) (Supplementary Table 3). Of note, we found the unsupervised clustering of the synthetic tau PET data separated the majority of patients with a Visuospatial clinical phenotype from those with a Language clinical phenotype. Taken together this indicates that the synthetic tau PET can capture phenotypically meaningful structure related to atypical clinical AD variants.

## Discussion

Here we introduce CoMA-UNet, a novel approach to incorporate non-PET covariate information in an MRI to tau-PET image synthesis. We show that the synthetic images produced using the CoMA-UNet were of high fidelity and markedly outperformed existing MRI-to-PET models. Furthermore, we show that the CoMA-UNet produced synthetic images that generalized across cohorts and tracers while preserving cognition-linked signals sufficient to support downstream clinical tasks. Critically, we provide an in-depth validation of the synthetic tau-PET images derived out-of-sample to the gold standard neuropathological Braak staging when images are generated with and without plasma biomarkers for AD.

CoMA-UNet synthesizes full volumetric tau PET images from structural MRI by incorporating subject-level covariates from non-PET variables (e.g. demographics, genetics, plasma markers) into the decoder of its UNet component. Through covariate-parameterized prompts anatomical priors are combined with ROI-level tau estimates through auxiliary models, yielding anatomically coherent SUVR patterns and accommodating covariate missingness. Previous deep learning approaches for volumetric MRI-to-PET using classical U-Net variants^20–22^, transformer backbones^23–26^, generative-adversarial paradigms, or drift-diffusion omit explicit covariate conditioning, relying on pure image-to-image mappings^18,27–35^. Prediction of tau-PET from non-imaging inputs such as plasma biomarkers, or from coarse imaging features such as MRI-derived regional summaries has been increasingly explored^15–17^, yet these approaches typically produce region-level or categorical estimates rather than full-resolution voxel-wise maps. In contrast, CoMA-UNet directly generates spatially resolved tau-PET representations, enabling finer-grained correspondence with ground-truth signal. This yielded a clear benefit over alternate convolutional and transformer-based architectures achieving substantially higher regional associations with ground truth tau PET (average ROI correlation CoMA-UNet: r= 0.52, others: r<0.15). Moreover, CoMA-UNet showed strong generalizability in the external NACC cohort, with synthetic tau PET generated without plasma markers capturing variance in an unseen tracer while outperforming other comparable modelling approaches on the same task^18^. Together, these findings demonstrate the potential of CoMA-UNet to generate high fidelity synthetic tau PET from MRI. Performance of MRI to tau PET synthesis using CoMA-UNet also are in line with FDG-PET to tau PET synthesis^36^. However, the use of MRI as an input imaging modality over FDG-PET has several important benefits. First, FDG-PET is less routinely collected in comparison to structural MRI ^37,38^. Second, FDG-PET is invasive requiring intravenous radiotracer administration and radiation exposure^37,39^. Third, FDG-PET acquisition is longer requiring the patient to wait before the collection of imaging within a post-injection window. These practical constraints, together with higher cost ^37^ make MRI a more readily available input imaging modality for the image synthesis problem^38^. This is evidenced by the relative paucity of FDG-PET in comparison to MRI in the large cohort and clinical trial samples we analyzed here.

We show through a series of downstream tasks that the synthetic tau PET generated by CoMA-UNet preserves cognition-linked signals sufficient for diagnostic classification and capturing participant-level variance in cognition. Synthetic tau PET images showed strong voxel-wise association with MMSE in association cortices, reflecting established findings that neocortical tau best explains cognitive decline^3,40–42^. Regionally, this association was significantly stronger than that of regional MRI features with MMSE in held-out Aβ positive subjects, aligning with results suggesting that tau PET is more predictive of cognition than MRI^43–45^. Comparing correlations between model derived and ground truth temporal meta-ROI SUVR with MMSE showed no statistical difference. This benchmarks synthetic tau PET generated by CoMA-UNet against prior work showing the utility of tau PET in discriminating different levels of clinical impairment^10,46^.

We present a potential use of synthetic tau PET imaging to retrospectively re-stratify clinical trials based on heterogeneity of baseline tau when tau PET was not available. When re-stratifying Aβ -positive cognitively normal older participants from the A4 solanezumab trial^19,47^ into tertiles of synthetic temporal meta-ROI SUVR we revealed significant divergence in PACC trajectories, with participants in the highest tertile showing the steepest cognitive decline. This is of clinical relevance as the successful TRAILBLAZER trial for Donanemab showed when participants were stratified by baseline tau burden treatment benefits varied^48^. Together, this offers an interesting avenue to apply synthetic tau PET to other completed clinical trials for AD, re-stratifying based on heterogeneity in baseline tau-PET and reassessing treatment effects. In doing so, retrospective subgroup discovery, counterfactual enrichment that prioritizes intermediate tau burden, and hypothesis testing on how tau modulates effect sizes, can all be achieved using MRI-only datasets for future tau-informed trial design.

We provide the most compelling account of the biological validity of tau-PET surrogates to date showing a tight coupling to the underlying neuropathological staging of AD. In two highly heterogenous held-out cohorts with neuropathology, voxel-wise Braak-group contrasts (I/II vs III/IV; III/IV vs V/VI) reproduced the canonical early medial-temporal tau focus, with progressive spread to posterior cingulate and retrosplenial cortex, lateral temporal and parietal association areas, and ultimately broader neocortex^42,49^. This finding was robust to the covariate feature set available (i.e. with or without plasma in the synthesis) and the standardization of MRI imaging. Namely, in the out-of-sample NACC cohort consisting of over 475 scanning sessions with non-SCAN compliant MRIs we accurately recovered the underlying Braak staging assessed at autopsy. This provides strong evidence that the synthetic tau PET generated using the CoMA-UNet aligns with unobserved ground truth tau PET that shows antemortem-postmortem concordance and Braak stage-like regional progression^3,40,50,51^. Taken together, these results indicate that even under heterogeneous, lower-quality MRI and covariate inputs, CoMA-UNet captures anatomically coherent Braak-ordered tau distributions, supporting its use in large retrospective MRI repositories lacking tau PET or complete covariates.

Finally, we show that CoMA-UNet can be adapted to atypical presentations of AD using a relatively small phenotype-enriched SCAN sample. In the atypical SCAN cohort, models fine-tuned from the model trained on the full ADNI-A4 dataset in a five-fold framework achieved strong held-out agreement with ground-truth tau PET, despite the atypical clinical phenotypes and associated topographic heterogeneity of the cohort ^52,53^. Representative subject-level examples further illustrated this point, with synthetic tau PET preserving ROI-level hemispheric percent-ratio patterns in a language-predominant atypical AD case, in contrast to the more symmetric and less strongly lateralized pattern observed in a typical AD comparison case (Figure *7*). The fidelity of this topographic reproduction was observed as a high centroid correspondence in a paired analysis of atypical NACC patients with ground truth tau PET available, indicating the dominant structures in the true atypical tau PET signal were preserved in the synthetic tau PET signal. Furthermore, in the atypical non-SCAN cohort, unsupervised clustering of synthetic tau PET regional signal was significantly associated with clinical domain label, with clusters separating many visuospatial-predominant from language-predominant cases. Together, these findings indicate that CoMA-UNet is not restricted to the typical AD tau pattern learned from the primary ADNI-A4 cohort but can be adapted to domain-relevant atypical tau topographies when fine-tuned within an appropriate phenotype-enriched sample.

The synthetic tau PET generated through CoMA-UNet has several limitations. Relative to modern vision-based deep learning models such as Stable Diffusion^33^, CoMA-UNet contains significantly fewer trainable parameters. This lightweight base configuration of CoMA-UNet presented here makes it widely accessible and deployable, although limiting its maximal performance ^54–56^. The framework, however, naturally supports parameter scaling, as additional covariate-parameterized prompts can be introduced to enable the learning of a broader set of covariate-specific representations^57,58^. Such extensions are most appropriate for covariates that define subject strata whose expected ground-truth distributions differ fundamentally from those of other groups^59–62^. Thus, while the present implementation is constrained by parameter count, the CoMA-UNet architecture provides a conceptually simple and extensible pathway for scaling. Furthermore, our current implementation of CoMA-UNet was trained exclusively to synthesize FTP tau PET from MRI. Given that tau PET tracers exhibit distinct profiles of off-target binding and region–specific sensitivity to tau pathology^63–65^, extending the model to jointly learn across multiple tracers represents an important direction for future work.

We present and validate the CoMA-UNet pipeline for high fidelity MRI to tau-PET synthesis with varied non-PET covariate data available. This novel approach out-performs other deep learning approaches, producing biologically valid, and clinically relevant synthetic tau PET. Through the application of this pipeline, retrospective analyses to investigate regional tau PET pathology in MRI-only or partially missing datasets is possible. Through these synthetic tau PET images, new avenues for predictive modelling are available allowing the aggregation of legacy data resources with publicly available tau PET samples or the precision re-stratification of completed clinical trials.

## Methods

### Participants and Data

#### Datasets

Evaluations span several held-out datasets, arising from three distinct parent cohorts. Each dataset was used to assess different generalization settings, including internal held-out performance, missing-modality deployment, inter tracer validation, neuropathological concordance, and fidelity of atypical AD topographies. Cross-sectional and cross-validation cohorts were drawn from the Alzheimer’s Disease Neuroimaging Initiative (ADNI) and the A4 study, comprising ADNI participants across the cognitive spectrum (cognitively normal, mild cognitive impairment, and Alzheimer’s dementia) and cognitively normal A4 participants, each with available T1-weighted MRI and [^18^F]flortaucipir (FTP) tau PET imaging. The ADNI-A4 training dataset (n = 1250 image pairs) was randomly partitioned into five folds, balanced for meta-temporal tau burden and amyloid-β (Aβ) positivity. All analyses performed within this dataset used a standard five-fold procedure: for each fold, CoMA-UNet was trained on four folds and used to synthesize tau PET only for that fold’s held-out subjects. For external dataset analyses, CoMA-UNet was trained once on the entirety of the ADNI-A4 training set. The cross-sectional validation dataset (n = 200 image pairs) comprised participants from ADNI (n = 100) and A4 (n = 100). Separately, for post-mortem validation, we held out 99 distinct ADNI participants with autopsy-confirmed Braak stage data. An additional external A4 validation cohort included unseen A4 participants with available T1 MRI but no corresponding tau PET, used to test generalization to missing-modality settings and sensitivity for prospective cognitive decline. Plasma biomarkers were quantified using Fujirebio Lumipulse assays (Aβ42 /Aβ40 and p-tau217 in participants from the ADNI and A4 cohorts) and Quanterix Simoa assays (Aβ42 /Aβ40, p-tau217, neurofilament light [NfL], and glial fibrillary acidic protein [GFAP] in participants from the ADNI cohort). For further external validation, we included participants from the National Alzheimer’s Coordinated Center (NACC), encompassing both SCAN-compliant and non-SCAN-compliant MRI acquisitions. The NACC SCAN (n = 1409) cohort had paired tau PET using either [^18^F]flortaucipir (n = 958) or [^18^F]MK-6240 (n = 451), providing heterogeneity in tracer kinetics and quantification scales. The NACC non-SCAN cohort (n = 475) consisted of participants with autopsy-confirmed Braak staging but no PET imaging, enabling neuropathological validation. Plasma biomarkers were unavailable for NACC participants. The atypical SCAN, and atypical non-SCAN cohorts included NACC participants with AD as the primary etiologic diagnosis, atypical clinical presentations, and available MRI. Samples in the atypical SCAN cohort had paired tau PET imaging. The atypical cohorts comprised 452 total samples with single-domain non-amnestic, multi-domain amnestic, and multi-domain non-amnestic impairment profiles, including participants with predominant visuospatial, attention, executive, and language-domain involvement. Because these atypical phenotypes were not represented in the primary ADNI-A4 training cohort, this analysis was treated separately from the fixed-model external validation analyses. Specifically, CoMA-UNet was evaluated using a dedicated five-fold cross-validation framework within the atypical SCAN sample: for each fold, the model trained on the full ADNI-A4 dataset was fine-tuned on four folds and used to synthesize tau PET only for the held-out fold. Held-out predictions were then stacked across folds to obtain out-of-sample synthetic tau PET images for all Atypical SCAN participants. To supplement downstream analyses, 99 subjects with available MRI-FTP tau PET image pairs were sampled from the SCAN NACC dataset. These subjects were sampled such that the distribution of age would match that of the atypical cohorts, and to achieve a balanced distribution of Sex.

SCAN compliance indicates that MRI data were prospectively acquired under NACC’s standardized SCAN protocol using approved acquisition parameters and centralized QC/harmonization procedures. In practice, this includes using approved acquisition parameters under the SCAN MRI protocol, collecting images prospectively within the SCAN framework, and passing the centralized SCAN processing/QC pipeline; images collected before 2021 or acquired under mixed/non-approved protocols are not considered SCAN-compliant, even though they may still be accepted by NACC as non-standard imaging data. As a result, the non-SCAN subset likely contains greater heterogeneity in acquisition characteristics and protocol adherence, which can introduce additional technical variance and make it a more challenging but also more realistic test bed for evaluating model generalization under less harmonized real-world conditions

#### Covariates & Feature Sets

The synthesis of the tau PET images from MRI proceeded in three stages, each using a distinct set of covariates. In the first stage, CatBoost models were used to estimate regional tau burden, MetaTempTau, Aβ status, whereas MMSE was estimated using a KNN model. These predictors use demographic (age, sex, education, marital status, self-reported race, height, weight), vital sign (systolic and diastolic blood pressure, pulse, respiratory rate, temperature), genetic (APOE4 allele count), and plasma biomarkers (p-tau217, GFAP, NfL, Aβ40, and Aβ42), when available. This expanded covariate set was included on pragmatic predictive grounds, with the goal of capturing subject-level disease context and feature interactions that may improve estimation of tau-related outcomes. In the second stage, the deep learning model integrated these MetaTempTau, Aβ status, and MMSE estimates through a conditional decoder. In the third stage, the regional tau estimates are used in a final modulation step. Note that plasma biomarkers were not provided directly to CoMA-UNet. For model performance analyses and all downstream evaluation tasks, MetaTempTau and regional tau values were derived exclusively from synthetic tau PET images. We emphasize that during inference, Aβ status and MMSE are estimated at all model stages and are never used directly.

#### Regions of Interest

In our analyses we consider *R* = 32 regions of interest (ROI) taken from the Desikan-Killiany (DK) Atlas (Supplementary Table 1). In particular, these ROIs span the parietal, temporal, and include the Hippocampus and Amygdala. For each of the ROIs, we consider the left and right hemispheres separately. In addition to these regions, we also utilize a Meta ROI, denoted as *MetaTempTau*, consisting of the Amygdala, Inferior and Middle Temporal, Entorhinal, Parahippocampal, and Fusiform regions. The SUVR in the *MetaTempTau* is calculated as the weighted average of SUVR in each of its constituent regions.

### Models

#### Deep learning model for synthetic tau PET generation

A novel deep learning model, Covariate-modulated Attention UNet, was used to synthesize tau PET scans from MRIs. The architecture of the model consists of an Attention UNet with conditional up-sampling steps integrating subject-specific covariate data and estimated pathological markers. In particular, we replace all up-sampling Conv(⋅) operations in the Decoder with Conditional Convolution operators^66^, integrating age, sex, education, as well as estimated MMSE and *MetaTempTau*.

To further increase the inductive bias and enhance the predictive capability of CoMA-UNet, estimated regional average tau values obtained from CatBoost^67^ models are incorporated. The core idea is to modulate the initial predictions using a dynamic prompt that is both parameterized and tailored based on individual covariates and estimated regional tau load. Through the integration of these individual average tau values directly into the prediction process, the model becomes more sensitive to variations in disease pathology among different subjects. This modulation framework thus enables the model to capture subtle variations in the data associated with covariates and pathological states, leading to more accurate and individualized assessments and enhancing its predictive performance. The MRI provides subject-specific anatomy and atrophy patterning, while demographics, genetics, and plasma biomarkers encode subject-level disease risk. The prompt module therefore does not replace image information but modulates image decoding so that the synthesized tau distribution is conditioned on both spatial anatomy and non-spatial subject context.

Formally, given an input MRI *M*_*i*_, the CoMA-UNET produces an initial prediction *Z*_*i*_ ∈ ℝ^*p*1×*p*2×*p*3^ . This output is refined by computing the final prediction *P̂*_*i*_ as *P̂*_*i*_ =ReLU(Conv(*Z*_*i*_ ⊕ConvBlocks(*F̃* ^*ci*^ ⊕ *Z*_*i*_)), where ReLU(⋅) is the Rectified Linear Unit activation function, and ⊕ represents concatenation along the channel dimension. The function ConvBlocks(⋅) consists of a sequence of three convolutional operations, each followed by PReLU activation, dropout, and instance normalization, processing the concatenated features to extract higher-level representations. The modulated dynamic prompt is computed as *F̃* ^*ci*^ = *F* + ConvBlocks(*F*^*ci*^ ⊕ *Ŝ*_*i*_ ⊕ 𝛤̂_*i*_), which adjusts a base dynamic prompt *F* using three subject-specific components. The tensor 𝛤̂_*i*_ ∈ ℝ^*p*1×*p*2×*p*3^ contains CatBoost-estimated regional tau SUVR values *tâu*_*r*,*i*_ placed in voxels corresponding to brain region *r* ∈ *R*, and zeros elsewhere, thereby masking out non-informative background. The tensor *Ŝ*_*i*_ contains corresponding regional uncertainty estimates *σ̂*_*r*,*i*_ for each *tâu*_*r*,*i*_, assigned to the same voxels. The parameterized dynamic prompt *F*^*ci*^, selected based on the subject’s covariate *c*_*i*_. In this work, we set *c*_*i*_ to be the Aβ positivity status estimated by a CatBoost model. The convolution Conv(⋅) applies further processing to the combined features of 𝛤̂_*i*_ and *F̃* ^*ci*^, through a convolutional layer followed by PReLU activation, dropout, and instance normalization. This formulation allows the model to incorporate individual pathological markers and covariate information directly into the feature space, enabling it to adjust its predictions based on personalized disease profiles and improving its ability to capture individual differences for enhanced prediction accuracy.

The CoMA-UNet was trained by minimizing a ROI-weighted MSE, *L*_*Gen*_, in which the squared residuals in voxels corresponding to ROI *r* are scaled by pre-specified non-zero weights, *w*_*r*_. This loss function measures the difference between the ground-truth tau PET and the generated synthetic tau PET; when *L*_*Gen*_ = 0, the synthetic tau PET and the actual tau PET are the same. Concretely, over a set of *n* MRI and actual tau PET pairs, 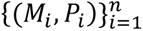, we define 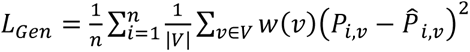, where *P̂* is the value of voxel *v* in the synthetic tau PET generated from *M*_*i*_, *V* is the set of all voxel coordinates, and *w*(*v*) is a function which either assigns a user-specified value of *w*_*r*_ ∈ ℝ if the voxel *v* is in ROI *r*, or a value of 1 otherwise.

#### Prediction of covariates and tau values

Missing covariate data were imputed using a KNN algorithm, with the number of neighbors equal to the square root of the total number of features, and each neighbor weighted uniformly. Independent CatBoost predictors were trained using plasma biomarker, family history, demographic, vital, and MRI-derived regional summary features to estimate temporal meta-ROI SUVR (*MetaTempTau*), as well as regional tau SUVR estimates (*tâu*_*r*,*i*_) and their associated standard deviation (*σ̂*_*r*,*i*_), assuming normally distributed errors. Using the same feature set, additional CatBoost models were separately trained to predict Aβ status, and a KNN model was used to predict MMSE scores. CatBoost and KNN model training and evaluation employed the same training and testing partitions as the deep learning model.

#### Image Processing

All T1-weighted MRI volumes were preprocessed using FreeSurfer v7.1^68^, which performed skull stripping, intensity normalization, cortical surface reconstruction, and subcortical segmentation of the native-space anatomical images. This process generated subject-specific anatomical volumes and corresponding Desikan-Killiany parcellations, providing voxel-wise anatomical label maps for subsequent region-based sampling. Each intensity-normalized MRI was then resampled to the model’s fixed grid of 128×128×128 voxels (*p*_1_ = *p*_2_ = *p*_3_ = 128) using nearest-neighbor interpolation, symmetrically padded along the axial axis, and centrally cropped before a single channel dimension was added to match the network’s expected input. Tau PET images using either [¹⁸F]Flortaucipir (FTP) or [¹⁸F]MK-6240 (MK) radiotracers corresponding to the T1 MRIs were processed following the Berkeley PET Imaging Pipeline (B-PIP)^69^, which includes framewise motion correction, frame averaging, co-registration to the subject’s MRI, and normalization to the inferior cerebellar gray reference region for standardized uptake value ratio (SUVR) computation. For select downstream evaluations, specified in the Evaluation section, the model’s output volumes were spatially normalized to MNI space (157×189×156 voxels) using nonlinear registration in SPM12 and resampled with third-order B-spline interpolation.

### Metrics

Similarity between ground-truth tau PET and synthetic tau PET was assessed across cohorts using regional SUVR calculated from *R* ROIs in addition to full-image wide metrics. Pearson’s correlation, Mean Absolute Percentage Error (MAPE), and Mean Absolute Error (MAE) were calculated in each ROI, with MAPE and MAE also being calculated across the entire image. To further quantify the quality of the synthetic images we calculate the average of the regional Pearson correlations (Corr_AVG_) and the structural similarity index (SSIM). SSIM is a metric used to measure the similarity between two images. SSIM is designed to improve on traditional methods like peak signal-to-noise ratio (PSNR) and mean squared error (MSE), which have been shown to be inconsistent with human visual perception. Instead, SSIM considers changes in structural information, luminance, and contrast. The SSIM index is a decimal value between -1 and 1, where 1 indicates perfect similarity. It is calculated based on the comparison of three main components of the images: luminance (l), contrast (c), and structure (s). We give the precise mathematical definitions of each of the described performance metrics in the Supplementary.

### Training & Evaluation

#### Training

The model was trained on an Nvidia Tesla V100 (16G VRAM) GPU using PyTorch^70^. Optimization of the generative loss function *L*_*Gen*_ was performed using AdamW with an initial learning rate of 0.001, weight decay of 0.001, and a reduce-on-plateau learning rate scheduler, with batch sizes of two. Early stopping was performed after the model didn’t improve on a held-out validation subset for five consecutive epochs. Model parameters were first tuned using a five-fold cross-validation framework, in which each fold was trained exclusively on its training partition and evaluated on the held-out partition via strict out-of-fold predictions. Following hyperparameter selection, a final model was retrained on the full cross-validation dataset (n = 1250) using the same optimization procedure and was subsequently used for all downstream validation analyses.

All auxiliary predictors for ROI-level tau, Aβ status, and cognitive score were exclusively fit on the same training split as the deep learning model. That is, in cross-validation, each auxiliary model was re-fit within every outer fold in lockstep with the DL model; for cross-sectional analyses, models were fit on the full cross-sectional training set. Thus, all evaluations relied strictly on out-of-sample predictions from every model component.

#### Evaluation

During five-fold cross-validation over the 1,250 samples in the ADNI-A4 training dataset, the model was trained on four folds (80%) and tested exclusively on the held-out fold (20%). Model performance for each fold was then quantified using the similarity metrics described in the Supplementary Metrics Section, including SSIM, MAPE, MAE, ROI-focused variants of MAPE and MAE, as well as regional correlation and Corr_AVG_. We present the cross-validation performance of the CoMA-UNet as benchmarked on the ADNI-A4 training dataset against several other model with alternative architectures. The CoMA-UNet model was then trained on the entirety of ADNI-A4 training dataset and was retained for evaluation on the downstream tasks to assess the clinical utility of synthetic tau PET scans. We elaborate each downstream task in the subsequent sections.

#### AD Diagnosis Classification

We conducted pairwise classifications across cognitively normal (CN), mild cognitive impairment (MCI), and Alzheimer’s disease Dementia (Dem) groups in the ADNI samples within the ADNI-A4 training and cross-sectional validation datasets. Samples from A4 were excluded in this analysis to prevent the model from being biased against population differences. For these pairwise classification tasks, we trained logistic regression models using the synthetic tau PET uptake in the temporal meta-ROI (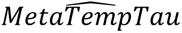). In the ADNI-A4 training dataset, we used a five-fold cross-validation scheme, computing 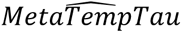 from the synthetic tau PET images generated from the MRIs in each held-out fold. Logistic-regression classifiers were trained on the remaining four folds and evaluated on the held-out fold. In the external cross-sectional validation dataset, 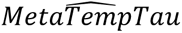 was classified using logistic-regression classifiers trained on the 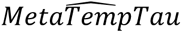 derived from synthetic tau PET images produced for held-out samples in the cross-validation. Model performance was assessed using receiver operating characteristic area under the curve (ROC-AUC) scores. To quantify voxel-level discriminative effect size across the brain, we computed voxel-wise Cohen’s d values in spatially normalized synthetic tau PET for each diagnostic cohort comparison.

#### Cognitive Baseline Prediction

To assess how well synthetic tau PETs relate to baseline cognition in comparison to actual tau PETs, we compared correlations of 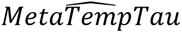-MMSE and *MetaTempTau*-MMSE with the cross-sectional cohort, quantifying this similarity using Steiger’s Z. We additionally evaluated the voxel-wise correlation between spatially normalized synthetic tau PET scans and MMSE. Finally, to assess incremental value over structural MRI, we extracted ROI-level SUVR and corresponding MRI regional volumes from the Desikan-Killiany atlas. For each ROI and Aβ stratum, we compared SUVR-MMSE versus MRI-MMSE correlations using Steiger’s Z for dependent correlations. Multiple comparisons across ROIs were controlled with the Benjamini-Hochberg False Discovery Rate procedure.

#### Braak Stage Contrasts

To test whether synthetic tau PET recapitulates the canonical neuropathological progression of tau, we performed adjacent-stage voxel-wise contrasts in the ADNI-NP cohort (n=99), and separately, in the NACC non-SCAN cohort (n = 475). Synthetic PET volumes were spatially normalized to MNI space and grouped according to Braak stages I/II, III/IV, and V/VI. We conducted two independent contrasts (I/II vs. III/IV; III/IV vs. V/VI) using two-sample independent t-tests at each voxel, and additionally computed Cohen’s d effect-size maps. As a complementary ROI analysis, we summarized medial-temporal (MTL) SUVR from each synthetic volume and fit an ordinal (proportional-odds) regression of MTL SUVR on ordered Braak group.

#### A4 population restratification

To examine whether model-derived measures could refine treatment group stratification, we first modeled PACC trajectories as a function of treatment assignment (placebo versus solanezumab) in the A4 validation cohort. We then re-stratified participants within each treatment arm according to 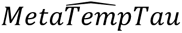 tertiles derived from synthetic tau PET scans, generated both with and without incorporation of plasma biomarker information to explore the impact of plasma-informed versus MRI-only synthetic PET stratifications. This approach enabled assessment of cognitive trajectories within more biologically informed subgroups, providing a framework to evaluate whether synthetic imaging markers could enhance sensitivity to treatment effects beyond conventional randomization strata. Trajectory estimation followed the same statistical framework used in the original A4 study. Specifically, a linear mixed-effects model with natural cubic splines was employed to capture nonlinear longitudinal change, estimated by restricted maximum likelihood^47^.

#### Evaluation under unseen radiotracer in held-out subject population

We performed a rigorous evaluation of the proposed model’s generalization by assessing performance in the NACC-SCAN cohort (n = 1409), which contained both tau PET acquired with FTP tracer (n = 958), and an unseen tracer, MK-6240 (n = 451). We did not provide the model with plasma biomarkers in this experiment. For each tracer and ROI, we computed Pearson’s r between predicted SUVR from synthetic PET and observed SUVR from the corresponding ground-truth PET, along with R^2^. Since FTP and MK differ in binding properties and quantitative scales, we restricted evaluation metrics to these scale-invariant, correlation-based measures.

#### Evaluation of atypical AD presentations

Unlike the external validation analyses, which used the fixed CoMA-UNet model trained on the full ADNI-A4 training dataset, the atypical SCAN analysis used a dedicated five-fold cross-validation framework. In each fold, the deep learning component of CoMA-UNet was initialized from the model trained on the full ADNI-A4 dataset and fine-tuned on four folds of the atypical SCAN cohort. In contrast, the auxiliary predictors were trained from scratch within each fold using only the corresponding four-fold atypical SCAN training partition. The fine-tuned CoMA-UNet and fold-specific auxiliary predictors were then evaluated only on the held-out fold. Held-out predictions were stacked across folds to generate out-of-sample synthetic tau PET images for all atypical SCAN participants. Inference only datasets not included in the held-out atypical SCAN folds, included the non-SCAN atypical and SCAN typical datasets, each of the five fold-trained models was applied independently to every participant, and the resulting synthetic tau PET volumes were averaged voxel-wise before ROI SUVR extraction.

Goodness-of-fit in the atypical SCAN cohort was assessed by comparing out-of-sample synthetic tau PET with corresponding ground-truth tau PET using Corr_AVG_, MAE, and SSIM. To evaluate whether synthetic tau PET preserved multivariate atypical tau topographies, we performed unsupervised K-NN clustering of regional SUVR profiles atypical datasets. Preprocessing was fit independently within each dataset, with no shared scaling or imputation parameters across cohorts. K-means clustering was applied to regional tau PET SUVR profiles. Analyses used 72 bilateral ROIs, including the amygdala, hippocampus, and Desikan–Killiany cortical regions, with the corpus callosum label excluded. Clustering was performed with k = 4 clusters using squared Euclidean distance, 100 random initializations, a maximum of 1000 iterations, and a fixed random seed. The same clustering pipeline was applied independently to SCAN ground-truth tau PET, SCAN cross-fold synthetic tau PET, and non-SCAN synthetic tau PET.

Clinical coherence of the resulting clusters was assessed by testing the association between NACC domain label and cluster assignment using Pearson’s chi-square test. Domain labels included executive, multi-domain, language, visuospatial, and attention phenotypes.

To assess structural reproducibility of cluster-level tau patterns across datasets, cluster centroids were computed as the mean raw, non-z-scored ROI SUVR vector within each cluster. Because cluster labels are arbitrary across independently fit clustering solutions, matched by maximizing pairwise Pearson centroid correlations. Matched centroid correlations quantified how closely tau topography templates identified in SCAN ground-truth PET were reproduced in non-SCAN synthetic PET. As a within-cohort reference, the same centroid-alignment procedure was applied between SCAN ground-truth and SCAN cross-fold synthetic tau PET.

## Data and Code Availability

Python scripts, model checkpoints, and help files are available on GitHub. NACC data can be requested and downloaded at https://naccdata.org. Data from A4 are available to download on the LONI website at https://ida.loni.usc.edu. Data used in the preparation of this article were obtained from the Alzheimer’s Disease Neuroimaging Initiative (ADNI) database (adni.loni.usc.edu). The ADNI was launched in 2003 as a public-private partnership, led by Principal Investigator Michael W. Weiner, MD. The primary goal of ADNI has been to test whether serial magnetic resonance imaging MRI), positron emission tomography (PET), other biological markers, and clinical and neuropsychological assessment can be combined to measure the progression of mild cognitive impairment (MCI) and early Alzheimer’s disease (AD). For up-to-date information, see www.adni-info.org.

## Acknowledgements

J.G. is supported by the Alzheimer’s Association (23AARF-1026883). Data collection and sharing for this project was funded by the Alzheimer’s Disease Neuroimaging Initiative (ADNI) (National Institutes of Health Grant U01 AG024904) and DOD ADNI (Department of Defense award number W81XWH-12-2-0012). ADNI is funded by the National Institute on Aging, the National Institute of Biomedical Imaging and Bioengineering, and through generous contributions from the following: AbbVie, Alzheimer’s Association; Alzheimer’s Drug Discovery Foundation; Araclon Biotech; BioClinica, Inc.; Biogen; Bristol-Myers Squibb Company; CereSpir, Inc.; Cogstate; Eisai Inc.; Elan Pharmaceuticals, Inc.; Eli Lilly and Company; EuroImmun; F. Hoffmann-La Roche Ltd and its affiliated company Genentech, Inc.; Fujirebio; GE Healthcare; IXICO Ltd.; Janssen Alzheimer Immunotherapy Research & Development, LLC.; Johnson & Johnson Pharmaceutical Research & Development LLC.; Lumosity; Lundbeck; Merck & Co., Inc.; Meso Scale Diagnostics, LLC.; NeuroRx Research; Neurotrack Technologies; Novartis Pharmaceuticals Corporation; Pfizer Inc.; Piramal Imaging; Servier; Takeda Pharmaceutical Company; and Transition Therapeutics. The Canadian Institutes of Health Research is providing funds to support ADNI clinical sites in Canada. Private sector contributions are facilitated by the Foundation for the National Institutes of Health (www.fnih.org). The grantee organization is the Northern California Institute for Research and Education, and the study is coordinated by the Alzheimer’s Therapeutic Research Institute at the University of Southern California. ADNI data are disseminated by the Laboratory for Neuro Imaging at the University of Southern California. The A4 Study was a secondary prevention trial in preclinical Alzheimer’s disease, aiming to slow cognitive decline associated with brain amyloid accumulation in clinically normal older individuals. The A4 Study was funded by a public-private-philanthropic partnership, including funding from the National Institutes of Health-National Institute on Aging, Eli Lilly and Company, Alzheimer’s Association, Accelerating Medicines Partnership, GHR Foundation, an anonymous foundation, and additional private donors, with in-kind support from Avid Radiopharmaceuticals, Cogstate, Albert Einstein College of Medicine and the Foundation for Neurologic Diseases. The companion observational Longitudinal Evaluation of Amyloid Risk and Neurodegeneration (LEARN) Study was funded by the Alzheimer’s Association and GHR Foundation. The A4 and LEARN Studies were led by Dr. Reisa Sperling at Brigham and Women’s Hospital, Harvard Medical School, and Dr. Paul Aisen at the Alzheimer’s Therapeutic Research Institute (ATRI) at the University of Southern California. The A4 and LEARN Studies were coordinated by ATRI at the University of Southern California, and the data are made available under the auspices of Alzheimer’s Clinical Trial Consortium through the Global Research & Imaging Platform (GRIP). The complete A4 Study Team list is available on: https://www.actcinfo.org/a4-study-team-lists/. We would like to acknowledge the dedication of the study participants and their study partners who made the A4 and LEARN Studies possible. The NACC database is funded by NIA/NIH Grant U24 AG072122. NACC data are contributed by the NIA-funded ADRCs: P30 AG062429 (PI James Brewer, MD, PhD), P30 AG066468 (PI Oscar Lopez, MD), P30 AG062421 (PI Bradley Hyman, MD, PhD), P30 AG066509 (PI Thomas Grabowski, MD), P30 AG066514 (PI Mary Sano, PhD), P30 AG066530 (PI Helena Chui, MD), P30 AG066507 (PI Marilyn Albert, PhD), P30 AG066444 (PI John Morris, MD), P30 AG066518 (PI Jeffrey Kaye, MD), P30 AG066512 (PI Thomas Wisniewski, MD), P30 AG066462 (PI Scott Small, MD), P30 AG072979 (PI David Wolk, MD), P30 AG072972 (PI Charles DeCarli, MD), P30 AG072976 (PI Andrew Saykin, PsyD), P30 AG072975 (PI David Bennett, MD), P30 AG072978 (PI Neil Kowall, MD), P30 AG072977 (PI Robert Vassar, PhD), P30 AG066519 (PI Frank LaFerla, PhD), P30 AG062677 (PI Ronald Petersen, MD, PhD), P30 AG079280 (PI Eric Reiman, MD), P30 AG062422 (PI Gil Rabinovici, MD), P30 AG066511 (PI Allan Levey, MD, PhD), P30 AG072946 (PI Linda Van Eldik, PhD), P30 AG062715 (PI Sanjay Asthana, MD, FRCP), P30 AG072973 (PI Russell Swerdlow, MD), P30 AG066506 (PI Todd Golde, MD, PhD), P30 AG066508 (PI Stephen Strittmatter, MD, PhD), P30 AG066515 (PI Victor Henderson, MD, MS), P30 AG072947 (PI Suzanne Craft, PhD), P30 AG072931 (PI Henry Paulson, MD, PhD), P30 AG066546 (PI Sudha Seshadri, MD), P20 AG068024 (PI Erik Roberson, MD, PhD), P20 AG068053 (PI Justin Miller, PhD), P20 AG068077 (PI Gary Rosenberg, MD), P20 AG068082 (PI Angela Jefferson, PhD), P30 AG072958 (PI Heather Whitson, MD), P30 AG072959 (PI James Leverenz, MD). The NACC database is funded by NIA/NIH Grant U24 AG072122. SCAN is a multi-institutional project that was funded as a U24 grant (AG067418) by the National Institute on Aging in May 2020. Data collected by SCAN and shared by NACC are contributed by the NIA-funded ADRCs as follows:

Arizona Alzheimer’s Center - P30 AG072980 (PI: Eric Reiman, MD); R01 AG069453 (PI: Eric Reiman (contact), MD); P30 AG019610 (PI: Eric Reiman, MD); and the State of Arizona which provided additional funding supporting our center; Boston University - P30 AG013846 (PI Neil Kowall MD); Cleveland ADRC - P30 AG062428 (James Leverenz, MD); Cleveland Clinic, Las Vegas – P20AG068053; Columbia - P50 AG008702 (PI Scott Small MD); Duke/UNC ADRC – P30 AG072958; Emory University - P30AG066511 (PI Levey Allan, MD, PhD); Indiana University R01 AG19771 (PI Andrew Saykin, PsyD); P30 AG10133 (PI Andrew Saykin, PsyD); P30 AG072976 (PI Andrew Saykin, PsyD); R01 AG061788 (PI Shannon Risacher, PhD); R01 AG053993 (PI Yu-Chien Wu, MD, PhD); U01 AG057195 (PI Liana Apostolova, MD); U19 AG063911 (PI Bradley Boeve, MD); and the Indiana University Department of Radiology and Imaging Sciences; Johns Hopkins - P30 AG066507 (PI Marilyn Albert, Phd.); Mayo Clinic - P50 AG016574 (PI Ronald Petersen MD PhD); Mount Sinai - P30 AG066514 (PI Mary Sano, PhD); R01 AG054110 (PI Trey Hedden, PhD); R01 AG053509 (PI Trey Hedden, PhD); New York University - P30AG066512-01S2 (PI Thomas Wisniewski, MD); R01AG056031 (PI Ricardo Osorio, MD); R01AG056531 (PIs Ricardo Osorio, MD; Girardin Jean-Louis, PhD); Northwestern University - P30 AG013854 (PI Robert Vassar PhD); R01 AG045571 (PI Emily Rogalski, PhD); R56 AG045571, (PI Emily Rogalski, PhD); R01 AG067781, (PI Emily Rogalski, PhD); U19 AG073153, (PI Emily Rogalski, PhD); R01 DC008552, (M.-Marsel Mesulam, MD); R01 AG077444, (PIs M.-Marsel Mesulam, MD, Emily Rogalski, PhD); R01 NS075075 (PI Emily Rogalski, PhD); R01 AG056258 (PI Emily Rogalski, PhD); Oregon Health & Science University - P30 AG066518 (PI Lisa Silbert, MD, MCR); Rush University - P30 AG010161 (PI David Bennett MD); Stanford – P30AG066515; P50 AG047366 (PI Victor Henderson MD MS); University of Alabama, Birmingham – P20; University of California, Davis - P30 AG10129 (PI Charles DeCarli, MD); P30 AG072972 (PI Charles DeCarli, MD); University of California, Irvine - P50 AG016573 (PI Frank LaFerla PhD); University of California, San Diego - P30AG062429 (PI James Brewer, MD, PhD); University of California, San Francisco - P30 AG062422 (Rabinovici, Gil D., MD); University of Kansas - P30 AG035982 (Russell Swerdlow, MD); University of Kentucky - P30 AG028283-15S1 (PIs Linda Van Eldik, PhD and Brian Gold, PhD); University of Michigan ADRC P30AG053760 (PI Henry Paulson, MD, PhD) P30AG072931 (PI Henry Paulson, MD, PhD) Cure Alzheimer’s Fund 200775 - (PI Henry Paulson, MD, PhD) U19 NS120384 (PI Charles DeCarli, MD, University of Michigan Site PI Henry Paulson, MD, PhD) R01 AG068338 (MPI Bruno Giordani, PhD, Carol Persad, PhD, Yi Murphey, PhD) S10OD026738-01 (PI Douglas Noll, PhD) R01 AG058724 (PI Benjamin Hampstead, PhD) R35 AG072262 (PI Benjamin Hampstead, PhD) W81XWH2110743 (PI Benjamin Hampstead, PhD) R01 AG073235 (PI Nancy Chiaravalloti, University of Michigan Site PI Benjamin Hampstead, PhD) 1I01RX001534 (PI Benjamin Hampstead, PhD) IRX001381 (PI Benjamin Hampstead, PhD); University of New Mexico - P20 AG068077 (Gary Rosenberg, MD); University of Pennsylvania - State of PA project 2019NF4100087335 (PI David Wolk, MD); Rooney Family Research Fund (PI David Wolk, MD); R01 AG055005 (PI David Wolk, MD); University of Pittsburgh - P50 AG005133 (PI Oscar Lopez MD); University of Southern California - P50 AG005142 (PI Helena Chui MD); University of Washington - P50 AG005136 (PI Thomas Grabowski MD); University of Wisconsin - P50 AG033514 (PI Sanjay Asthana MD FRCP); Vanderbilt University – P20 AG068082; Wake Forest P30AG072947 (PI Suzanne Craft, PhD); Washington University, St. Louis - P01 AG03991 (PI John Morris MD); P01 AG026276 (PI John Morris MD); P20 MH071616 (PI Dan Marcus); P30 AG066444 (PI John Morris MD); P30 NS098577 (PI Dan Marcus); R01 AG021910 (PI Randy Buckner); R01 AG043434 (PI Catherine Roe); R01 EB009352 (PI Dan Marcus); UL1 TR000448 (PI Brad Evanoff); U24 RR021382 (PI Bruce Rosen); Avid Radiopharmaceuticals / Eli Lilly; Yale P50 AG047270 (PI Stephen Strittmatter MD PhD); R01AG052560 (MPI: Christopher van Dyck, MD; Richard Carson, PhD); R01AG062276 (PI: Christopher van Dyck, MD); 1Florida - P30AG066506-03 (PI Glenn Smith, PhD); P50 AG047266 (PI Todd Golde MD PhD)

