## Supplementary Text for "Neuropathologically validated MRI to tau PET synthesis via Covariate-modulated attention networks"

### Figures and Tables

| Parietal Lobe (left and right) | Temporal Lobe (left and right) | Medial Temporal (left and right) |
|---|---|---|
| Inferior Parietal, Superior Parietal, Supramarginal, Postcentral, Precuneus | Bankssts, Entorhinal, Fusiform, Superior Temporal, Middle Temporal, Inferior Temporal, Transverse Temporal, Temporal Pole, Parahippocampal | Hippocampus, Amygdala |

*Supplementary Table 1 Full list of regions which make up the R=32 regions of interest (ROI).*

| Model | MAE (std) | MAPE (std) | SSIM (std) | $Corr_{AVG}$ (std) | Mean ROI MAPE (std) |
|---|---|---|---|---|---|
| Attention UNET | 0.0050 (0.0015) | 3186.33 (548.14) | 0.7506 (0.0933) | 0.1024 (0.0394) | 13.05 (1.43) |
| UNET | 0.00888 (0.00249) | 2873.20 (709.58) | 0.5756 (0.2454) | 0.1108 (0.0436) | 24.82 (1.80) |
| UNETR | 0.0136 (0.0053) | 150753.1 (198027.5) | 0.644 (0.098) | 0.079 (0.033) | 13.07 (1.40) |
| Attn UNETR | 0.0159 (0.0004) | 112828.2 (74154.1) | 0.600 (0.017) | 0.096 (0.052) | 12.96 (2.21) |

*Supplementary Table 2 Cross-validation performance of alternative Deep Learning architectures in Mean Absolute Error (MAE), Mean Absolute Percentage Error (MAPE), Structural Similarity Index (SSIM), Average Region of Interest (ROI) Correlation ($Corr_{AVG}$.), and mean ROI MAPE. The Attention UNet model generally performs the best, achieving the highest SSIM, lowest MAE, second-lowest mean ROI MAPE, and second highest Average ROI correlation.*

| Domain label | Cluster 0 | Cluster 1 | Cluster 2 | Cluster 3 |
|---|---|---|---|---|
| Attention | 1.000 | 0.000 | 0.000 | 0.000 |
| Executive | 0.162 | 0.397 | 0.294 | 0.147 |
| Language | 0.209 | 0.256 | 0.535 | 0.000 |
| Multi | 0.224 | 0.289 | 0.355 | 0.132 |
| Visuospatial | 0.200 | 0.600 | 0.100 | 0.100 |

*Supplementary Table 3 Domain-label proportions by cluster for cross-fold predicted atypical non-SCAN synthetic tau PET.*

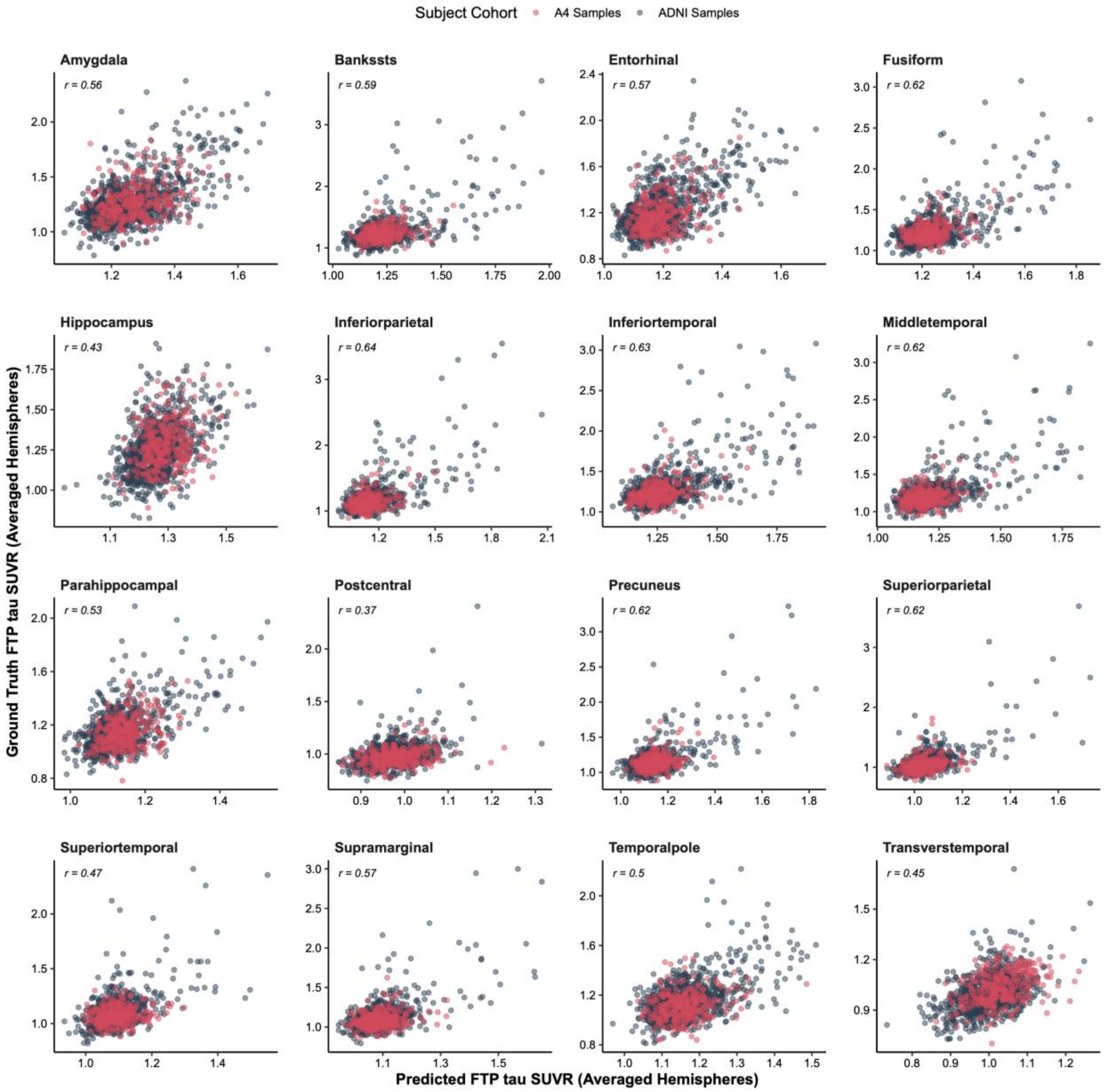


*Supplementary Figure 1 Hemisphere-averaged SUVR values for each ROI from ground-truth FTP tau PET versus synthetic FTP tau PET produced by Covariate-Modulated Attention UNet (CoMA-UNet), evaluated in the held-out test samples of each fold of the cross-validation splits of the ADNI-A4 training dataset. Samples from the A4-LEARN cohort are displayed with red markers, and samples from the ADNI dataset are displayed with dark-blue markers.*

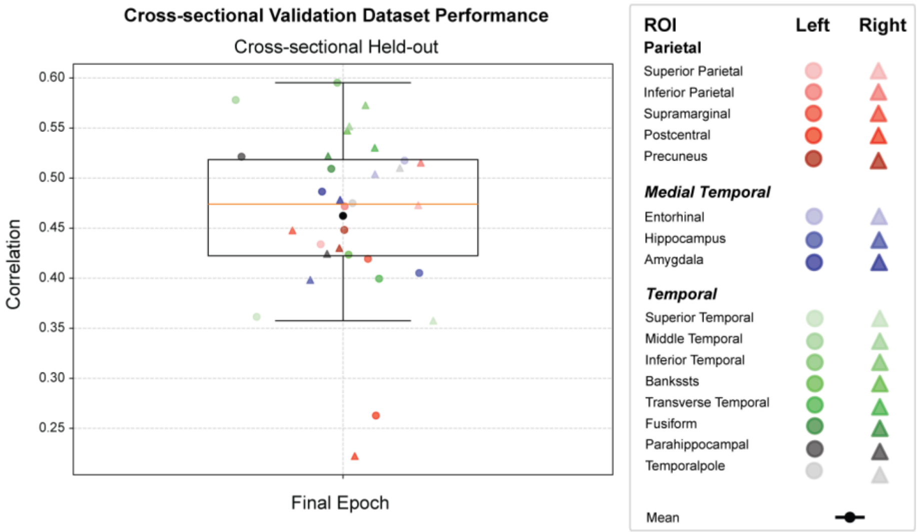


*Supplementary Figure 2 Cross-sectional validation dataset performance for the final trained model, showing ROI-level correlations between predicted and observed regional FTP SUVR in the independent test set. Higher correlation reflects more accurate regional tau estimation, with temporal and parietal regions showing the strongest predictive agreement, recapitulating cross-validation findings.*

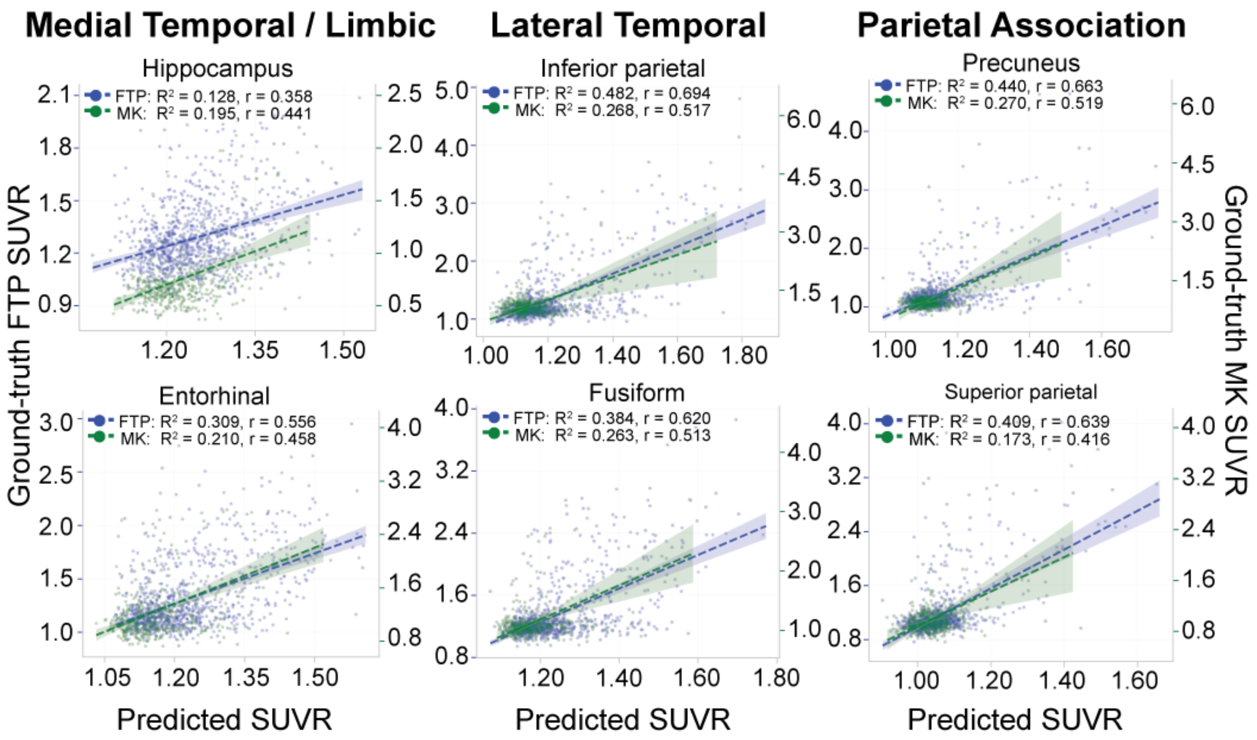


*Supplementary Figure 3 Correlations of synthetic tau SUVR with actual tau SUVR in key ROIs within the NACC SCAN dataset. Points corresponding to FTP tau SUVRs are displayed in blue, and points corresponding to MK-6240 tau SUVRs are displayed in green. The left-hand side y-axis marks the range of ground-truth FTP SUVRs, and the right-hand side y-axis marks the range of ground-truth MK-6240 SUVRs. Strong correlations are observed for both tracers.*

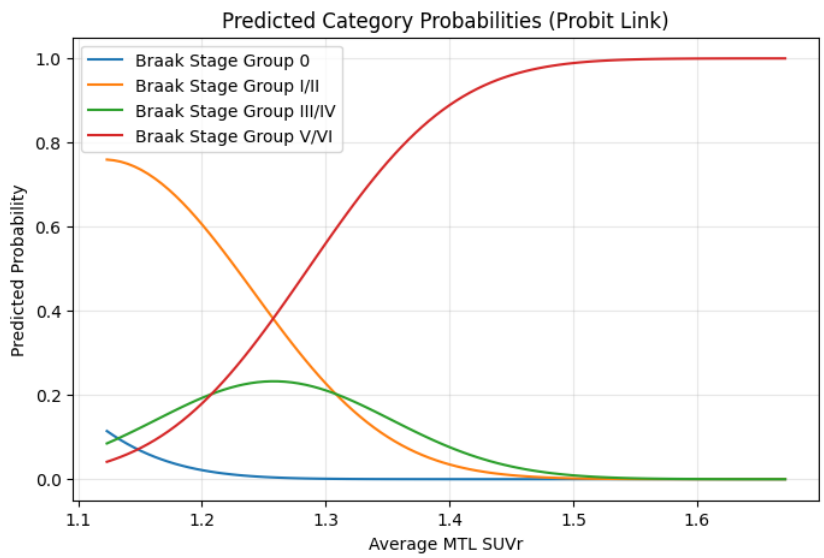


*Supplementary Figure 4 Ordinal regression on autopsy-confirmed Braak stage groups using average MTL SUVR from synthetic tau PETs within the ADNI-NP cohort.*

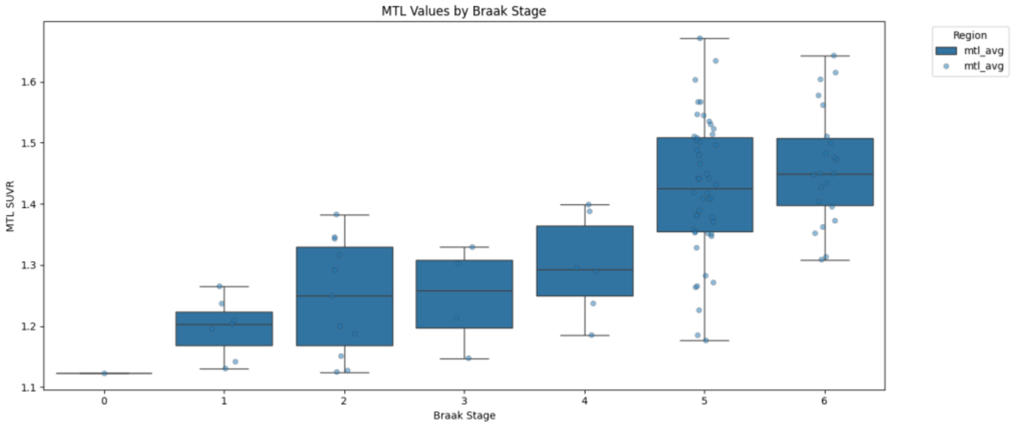


*Supplementary Figure 5 Average MTL SUVR by autopsy-confirmed Braak stage within the held out ADNI with available neuropath.*

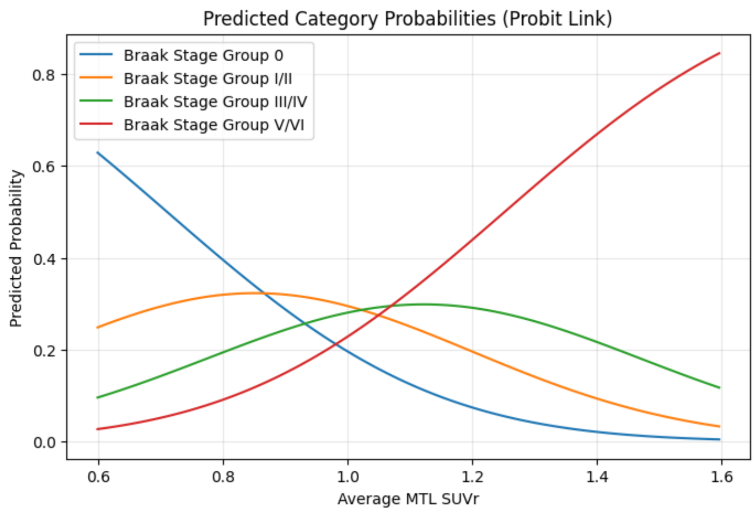


*Supplementary Figure 6 Ordinal regression on Braak stage group using average MTL SUVR from synthetic tau PETs within the NACC non-SCAN cohort.*

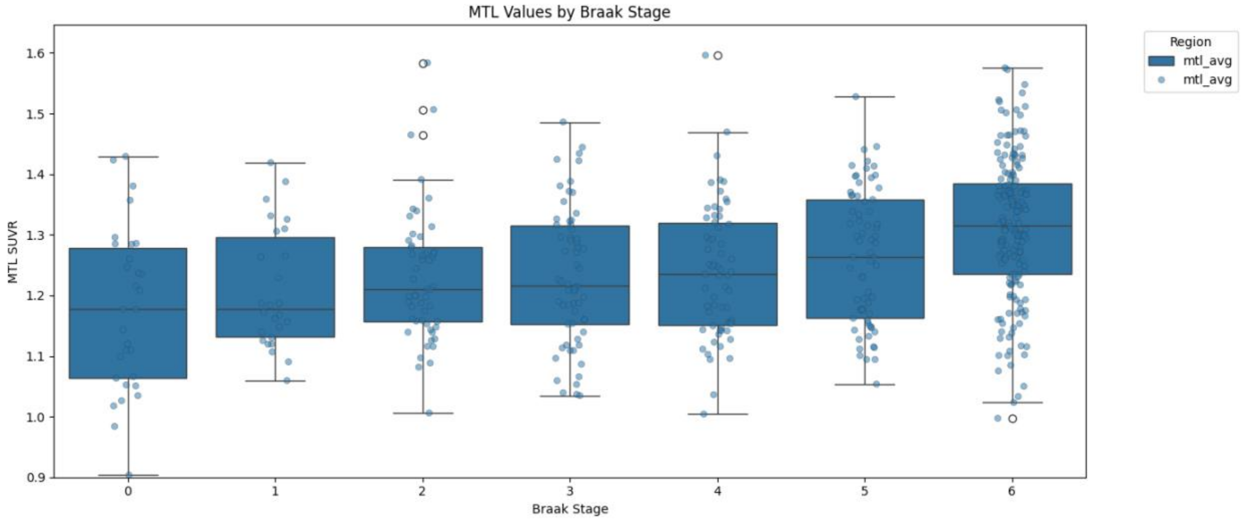


*Supplementary Figure 7 Average MTL SUVR by autopsy-confirmed Braak Stage within the NACC non-SCAN cohort.*

### Metrics

Within a cohort of patients indexed from $i = 1, \ldots, m$, let $\hat{P}_i$ be a synthetic tau PET generated from MRI $M_i$, and $P_i$ be the corresponding ground-truth tau PET. We define the following validation metrics:

- Mean Absolute Error (MAE). MAE of sample $j$ can be described as $MAE_i(\hat{P}_i, P_i) = \frac{||vec(\hat{P}_i - P_i)||_1}{p_1 \times p_2 \times p_3}$, where $vec(\cdot)$ is the vectorization operation to a tensor and $|| \cdot ||_1$ is the $L_1$ norm. The overall MAE across the subject cohort is $MAE = \frac{1}{n}\sum_{j=1}^{m} MAE_i(\hat{P}_{i,}P)$.
- Mean Absolute Percent Error (MAPE). MAPE of sample $j$ can be described as: $MAPE = ||vec(\hat{P}_i - P_i)/vec(P_i)||_1 \times 100\%$, where "/" is the element-wise division. The overall MAPE is thus defined as $MAPE = \frac{1}{m}\sum_{j=1}^{m} MAPE_i(\hat{P}_i, P_i)$.
- Region of Interest Correlation ($Corr_k$). We define an index matrix for a brain region of interest $R_1, \ldots, R_r \in \mathbb{R}^{p_1 \times p_2 \times p_3}$ with $r = 32$. For the $k$-th ROI and an arbitrary PET scan $P$ we define the mean SUVR as $SUVR_k(P) = \frac{||vec(P \odot R_k)||_1}{||R_k||_0}$. Then, for the $k$-th ROI across samples $j = 1, \ldots, m$, we generate two vectors: $V_{k,Pred} = \left(SUVR_k(\hat{P}_1), \ldots, SUVR_k(\hat{P}_m)\right)$ and $V_{k,True} = \left(SUVR_k(P_1), \ldots, SUVR_k(P_m)\right)$. Finally, the correlation for the $k$-th ROI is calculated as: $Corr_k = Corr(V_{k,Pred}, V_{k,True})$.
- Averaged ROI Correlation ($Corr_{AVG}$). Using the definition for ROI Correlation, we define $Corr_{AVG} = \frac{1}{r}\sum_{k=1}^{r} Corr_k$.
